# Loss of pod strings in common bean is associated with gene duplication, retrotransposon insertion, and overexpression of *PvIND*

**DOI:** 10.1101/2022.01.05.475151

**Authors:** Travis Parker, José Cetz, Lorenna Lopes de Sousa, Saarah Kuzay, Sassoum Lo, Talissa de Oliveira Floriani, Serah Njau, Esther Arunga, Jorge Duitama, Judy Jernstedt, James R. Myers, Victor Llaca, Alfredo Herrera-Estrella, Paul Gepts

## Abstract

Regulation of fruit development has been central in the evolution and domestication of flowering plants. In common bean (*Phaseolus vulgaris* L.), a major global staple crop, the two main economic categories are distinguished by differences in fiber deposition in pods: a) dry beans with fibrous and stringy pods; and b) stringless snap/green beans with reduced fiber deposition, but which frequently revert to the ancestral stringy state. To better understand control of this important trait, we first characterized developmental patterns of gene expression in four phenotypically diverse varieties. Then, using isogenic stringless/revertant pairs of six snap bean varieties, we identified strong overexpression of the common bean ortholog of *INDEHISCENT* (*PvIND*) in non-stringy types compared to their string-producing counterparts. Microscopy of these pairs indicates that *PvIND* overexpression is associated with overspecification of weak dehiscence zone cells throughout the entire pod vascular sheath. No differences in *PvIND* DNA methylation were correlated with pod string phenotype. Sequencing of a 500 kb region surrounding *PvIND* in the stringless snap bean cultivar Hystyle revealed that *PvIND* had been duplicated into two tandem repeats, and that a *Ty1-copia* retrotransposon was inserted between these tandem repeats, possibly driving *PvIND* overexpression. Further sequencing of stringless/revertant isogenic pairs and diverse materials indicated that these sequence features had been uniformly lost in revertant types and were strongly predictive of pod phenotype, supporting their role in *PvIND* overexpression and pod string phenotype.

**Significance:** Fruit dehiscence is a key trait for seed dissemination. In legumes, e.g., common bean, dehiscence relies on the presence of fibers, including pod “strings”. Selections during domestication and improvement have decreased (dry beans) or eliminated (snap beans) fibers, but reversion to the fibrous state occurs frequently in snap beans. In this article, we document that fiber loss or gain is controlled by structural changes at the *PvIND* locus, a homolog of the Arabidopsis *INDEHISCENT* gene. These changes include a duplication of the locus and insertion/deletion of a retrotransposon, which are associated with significant changes in *PvIND* expression. Our findings shed light on the molecular basis of unstable mutations and provide potential solutions to an important pod quality issue.

**Competing Interest Statement:** The authors have no competing interests.

## Introduction

Novel forms of fruit-mediated seed dispersal have been important to the evolutionary success of flowering plants. These depend on unique developmental programs which have evolved across taxa. In the Fabaceae, the third most speciose plant family (LPWG 2017), seed dispersal is typically mediated by explosive pod dehiscence, or shattering. For pod shattering to occur, multiple lignified pod layers must develop properly, including vascular sheath fibers or pod “strings”, and this is determined by a network of transcription factors and downstream cell wall modifying genes (Parker et al. 2021a).

Similar to legumes, *Arabidopsis thaliana* produces dehiscent seed pods, called siliques, whose development is under the control of a network of transcription factors (Di Vittori et al. 2019, Parker et al. 2021a). Among these, *INDEHISCENT (IND), SHATTERPROOF1/2*, and *ALCATRAZ* specify the valve margin region along which dehiscence occurs, and their expression is spatially restricted by genes such as *REPLUMLESS* and *FRUITFUL*. These patterning genes ultimately promote secondary cell wall formation and lignification through downstream genes, particularly the NAC and MYB family transcription factors (Nakano et al. 2015, Ohtani and Demura 2019), which are known to play a role in legume pod dehiscence (e.g., Rau et al. 2019, Zhang and Singh 2020). Other legume shattering-related genes, such as the dirigent gene *PDH1* of soybean and *PvPdh1* of common bean, influence pod torsion without anatomical changes (Suzuki et al. 2009, Parker et al. 2020, 2021b) and are unlikely to regulate pod strings.

Members of the legume family have been independently domesticated at least 40 times (Hammer and Khoshbakht 2015). Of these, common bean (*Phaseolus vulgaris* L.) is the largest source of plant protein and micro-nutrition for direct human consumption (Parker and Gepts 2021). The species was first domesticated (Piperno 2012) for its protein-rich dry beans. In wild beans, strongly lignified fibers exist at the pod sutures and inside the pod walls. These fibers lead to explosive pod dehiscence (or “shattering”) at maturity and, hence, seed dispersal. Following domestication, pod fiber was significantly reduced, although not eliminated. The suture fibers, also known as vascular sheath fibers, remained strong enough to be removed as pod “strings”, but led to undesirable toughness and stringiness in green beans. In the 19th century, breeder Calvin Keeney (Barnes 2010) identified a stringless mutation in the cultivar ‘Refugee Wax’, leading to a novel commercial class, namely “snap” beans grown for edible pods (Wallace et al. 2018). Non-stringy “snap” beans have become the global market standard for types grown as vegetables.

The main characteristic distinguishing dry and snap beans is the degree of fiber deposition in pods (Wallace et al. 2018). Non-stringy snap beans have lost secondary cell wall thickening of the vascular sheath, and lack a discernible dehiscence zone (Prakken 1934, Murgia et al. 2017, Rau et al. 2019, Parker et al. 2021a). Snap bean varieties display frequent spontaneous reversion to high pod fiber content (Fig. 1), for pod strings and/or pod wall fiber. Reversions that affect either trait individually or both simultaneously occur in all known snap bean varieties (Smith et al. 1997, Hagerty et al. 2016). Approximately 0.5-2.25% of plants in a population revert on average (Hagerty et al. 2016). This high-frequency reversion indicates that the trait may be controlled by transposable elements (Lisch 2013, Hirsch and Springer 2017), epigenetic factors such as DNA methylation (Miryeganeh and Saze 2020), or perhaps by rapidly evolving sequence repeats (Gemayel et al. 2012). This reversion also presentsa highly isogenic system to study the basis of pod string development and the mechanism of this high-frequency reversion.

**Fig 1.**
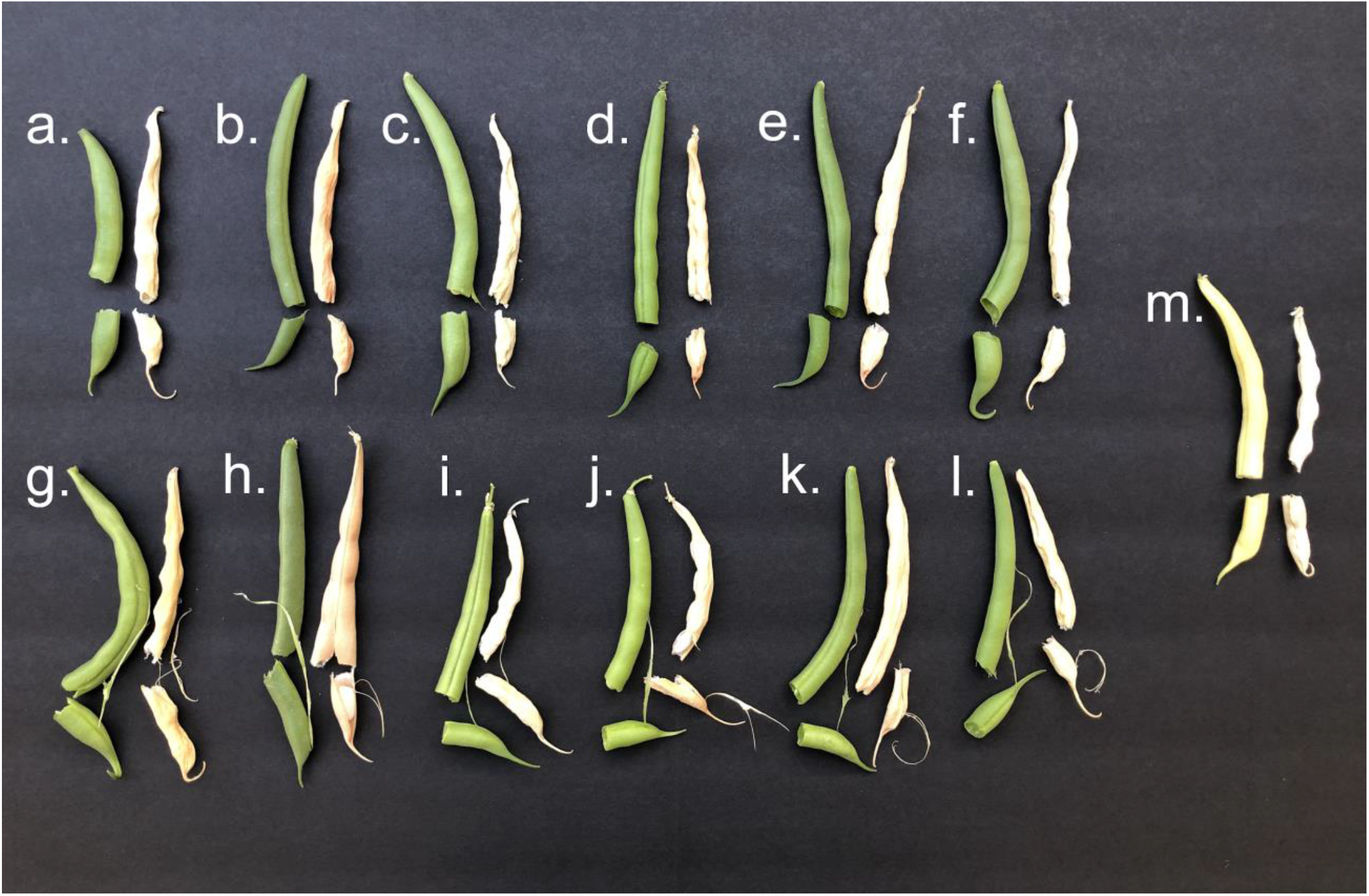
Pod phenotypes of stringless/stringy-revertant materials. a)-f) Six stringless cultivars of snap bean. No suture string is present in pods at seed fill stage (stage R8, Fernández et al. 1983) or dry mature pods. g)-l) Stringy revertants of each of the varieties shown above. When these pods are broken, a strong string can easily be removed from the sutures at both maturity stages. a) and g), cv. ‘Pismo’; b) and h), cv. ‘Prevail’; c) and i), cv. ‘Hystyle’; d) and j), cv. ‘Galveston’; e) and k), cv. ‘BBL156’; f) and l), cv. ‘Huntington’; m), stringless wax bean cv. ‘Midas’, which has been frequently included in studies of pod fiber traits (e.g. Koinange et al. 1996, Gioia et al. 2013, Murgia et al. 2017, Rau et al. 2019, Parker et al. 2020, Di Vittori 2021).

Emerson (1904) determined that pod strings were recessive, despite being the ancestral state, and only partly fit Mendelian segregation ratios. Drijfhout (1978) proposed that the dominant *String* (*St*) allele was required for any reduction in pod string, with a dominant hypostatic allele *Temperature Sensitive* (*Ts*) able to recover partial pod string in the presence of *St* at elevated temperatures. Koinange et al. (1996) mapped *St* to chromosome Pv02. In their analysis, pod wall fiber and pod suture fiber were genetically co-located, although other authors have found them unlinked (Emerson 1904, Prakken 1934, Hagerty et al. 2016). Gioia et al. (2013) described a *P. vulgaris* ortholog of *IND*, known as *PvIND*, which mapped 7.8cM from *St*. The occasional ‘recombination’ between *St* and *PvIND*, as well as a lack of explanatory genetic variation at the locus and 1 kb of promoter, suggested that this specific region was not responsible for control of pod strings. Hagerty et al. (2016) subsequently identified flanking markers for *St* spanning from approximately 500 kb from 43,984,700 to 44,472,300 (G19833 genome v2.1). This region includes four plausible candidate genes for the control of pod strings, namely two NAC transcription factors, one MYB family transcription factor, and *PvIND*.

Here, we investigate the genetic and transcriptional control of pod string development in common bean. To this end, we compare transcriptional patterns of diverse genotypes, including stringless/revertant pairs, identify anatomical effects of differentially expressed candidate genes, screen the methylation state of select regions of interest, and explore sequence variation in candidate regions across *P. vulgaris*.

## Materials and Methods

### Plant Materials

RNA-seq was conducted on four genotypes (G12873, ICA Bunsi, SXB 405, and Midas), which span the full range of pod fiber and shattering properties found in common bean (Supplemental Table S1). For RT-qPCR, anatomical studies, and bisulfite sequencing, eight pairs of stringy revertant pods and non-revertant stringless controls were collected at Syngenta facilities in 2017 (Supplemental Table S2). These were subsequently planted in the greenhouse in Davis, CA for pod sampling (Fig. 1). All seeds bred true to type for pod and other traits, without further reversion or instability in subsequent generations. One non-revertant snap type of the accession Hystyle was used for sequencing the full region between the flanking markers of the pod string locus of Pv02.

A set of 100 diverse *P. vulgaris* accessions were acquired from NPGS, UC Davis, and Oregon State University, and were greenhouse-grown in Davis, CA. DNA was extracted from leaf material or greenhouse-grown seeds using a modified CTAB method (based on Allen et al. 2006). Mature full-sized pods of each type were allowed to dry and were broken to analyze pod string phenotype on a scale of 0 (no removable string) to 10 (string readily removable).

### Pod string candidate and control genes

The 56 gene models between *St* flanking markers (Phvul.002G269200 to Phvul.002G274700; Hagerty et al. 2016) were accessed via the Legume Information System (Dash 2016; https://legumeinfo.org/home). Gene family and GO term data were downloaded through PhytoMine (Goodstein et al. 2012). Two NAC family transcription factors (‘*NAC 1*’, Phvul.002G271700; and ‘*NAC 2*’, Phvul.002G273100), one MYB family transcription factor (‘*MYB*’, Phvul.002G269900), and an atypical bHLH transcription factor closely related to *IND* (*PvIND*, Phvul.002G271000) were identified in the region. These were considered candidate genes for RNA-seq and RT-qPCR. For RT-qPCR, *Act11* (Phvul.008G011000) and *Ukn1* (Phvul.011G023200) were used as stably-expressed reference gene controls based on Borges et al. (2012). Amino acid sequences of *IND* homologs in *P. vulgaris* and Arabidopsis were compared using a fast minimum evolution tree based on the Grishin protein matrix on the NCBI website (blast.ncbi.nlm.nih.gov).

### RNA-seq

Gene annotation and GO data of the *Phaseolus vulgaris* v2.1 genome were downloaded from Phytozome (http://phytozome.jgi.doe.gov/). TopGO v2.26.0 was used to determine GO term enrichments. Transcriptomes were characterized using three pod replicates with pairwise comparisons between each of three different stages (Fernández et al. 1983): pod formation (R7), pod fill (R8) and pod maturation (R9). RNA sequencing libraries were prepared following the Illumina® TruSeq® Stranded Total RNA Sample Preparation kit instructions. A total of 36 TruSeq libraries were sequenced using the Illumina NextSeq500 platform in the 1×75 single-end mode, obtaining an average of 11.4 million raw reads per library. Raw RNA-seq data were processed with FastQC Version 0.11.2. Sequences with QC below 20 were trimmed using TRIMMOMATIC (Bolger et al. 2014) and adapters and overrepresented sequences were eliminated, obtaining about 10.8 million high-quality reads per library. The resulting reads of the good quality libraries were mapped with KALLISTO to *P. vulgaris* v2.1 from Phytozome. DEGs were determined with edgeR (v 3.16.5; Robinson et al. 2010) in R (v 3.3.2; R Core Team 2013) using a two-fold change threshold and FDR < 0.05.

To identify transcripts that were differentially expressed between stringy and non-stringy accessions, the expression patterns of ICA Bunsi, SXB 405, and G12873 were compared against those of Midas to establish GO functional category enrichments. Expression of candidate loci was also compared by ANOVA of linear model using genotype and maturity stage as fixed variables.

### RT-qPCR

Pods were harvested for RT-qPCR at 5 and 21 days after flowering (DAF). All samples from each developmental stage were harvested at the same time and date. Whole pods were harvested at five DAF; at 21 DAF, pod cross sections 1 cm thick were sampled at the first seed, with seed material immediately removed. Samples were flash-frozen in liquid N_2_ then kept frozen at −80°C. For RNA extraction, pods were ground in liquid N_2_ using a mortar and pestle, and extracted with the RNeasy Plant minikit (Qiagen, Hilden Germany). Concentration and quality were checked by NanoDrop and bleach gel. 500μg of RNA were used per sample for cDNA synthesis. Reverse transcription was conducted with the SuperScript IV VILO Master Mix with ezDNase kit (Invitrogen, Waltham MA USA). Transcript identifiers were used to generate primers for 70-150 bp amplicons using NCBI Primer BLAST (Supplemental Table S3), with specificity checking enabled to avoid amplifying non-target transcripts. Intron-spanning primers were used for multi-exon genes (‘*NAC 1’, ‘NAC 2’, Act11, Ukn1*). This was not possible for the single-exon genes ‘*MYB*’ and *PvIND*, so a control not treated with reverse transcriptase was included for these. qPCR primer efficiency was checked on pooled cDNA from each pod harvest date. All primers performed with an efficiency greater than >1.00. C_t_ values of reference genes were subtracted from those of candidate genes to generate ΔC_t_ data. These values are logarithmically related to RNA quantity, so they were converted to 2^(-ΔC_t_) values. The mean, standard deviation, and standard errors of 2^(-ΔC_t_) data for each phenotypic class was calculated for each gene comparison. Expression differences between the stringy and non-stringy groups were then compared by *t*-test.

### Microscopy

Full-sized green pods (stage R8) were harvested from the stringless/revertant pairs used for RT-qPCR, and 100μm transverse sections were made using a Vibratome. These were treated with Auramine O (0.01%) and Calcofluor (0.007%) for 20 min (Lo et al. 2021). Fluorescence was visualized using an Olympus BH2-RFL microscope (Waltham, MA, USA) with the ultraviolet filter set (UG-1 and DM-400 + L-420).

### Conserved *PvIND* promoter motifs

Upstream regulatory sequences of *PvIND* orthologs were compared to identify conserved elements with a potential role in transcriptional regulation. The *PvIND* amino acid sequence was retrieved from Phytozome 13 (phytozome-next.jgi.doe.gov). and pBLAST was used to find highly similar proteins in Arabidopsis and all legume species with available proteomes. DNA sequences upstream of these genes were downloaded and aligned with 2500 bp of the comparable *PvIND* region using NCBI BLASTn. In total, sequences upstream of 21 gene models representing 16 species were compared. The *PvIND* promoter was screened for enhancer activity using EnhancerPred (Jia and He 2016). *PvIND* homologs in common bean and Arabidopsis were also compared using NCBI BLAST tree viewer to compare relationships between these related proteins.

### Bisulfite sequencing

Bisulfite sequencing was conducted to analyze DNA methylation patterns potentially related to reversible phenotypic change. These areas included three conserved motifs upstream of *PvIND*, the area surrounding the *PvIND* start site, a region within the *PvIND* gene body, and part of the 3’ UTR (Supplemental Table S4). Genomic DNA was extracted using a modified CTAB protocol. Bisulfite treatment was conducted with the EZ DNA Methylation-Lightning Kit. Primers were designed with Zymo Bisulfite Primer Seeker 12S (Supplemental Table S4), PCR products were checked on a gel, cleaned with a QIAquick PCR Purification Kit, and genotyped by Sanger sequencing at the UC Davis DNA sequencing facility. FASTQ reads were converted to FASTA and aligned using NCBI BLAST to identify sequence variation. Methylation status of all cytosines was compared between non-stringy and revertant types.

### Sequencing 500 kb surrounding *PvIND*

To systematically search for variation that might cause differential expression of *PvIND*, the entire region between the *St* flanking markers (Hagerty et al. 2016) was sequenced and scaffolded in the stringless cultivar ‘Hystyle’ (sample Nampa-5) by Corteva Agrisciences (Johnston, IA). DLS BioNano mapping was combined with HiFi PacBio sequencing to create a single hybrid scaffold spanning the region. The scaffold was then aligned with the seven *Phaseolus* reference genomes of the genus *Phaseolus* on Phytozome to identify unique sequence features. Structural variation was evaluated using a panel of 100 stringy and non-stringy beans. PCR primers were developed using NCBI-BLAST (Supplemental Table S5) to span A) the retrotransposon site and B) the tandem duplication splice site, which also included the retrotransposon insertion. PCR was conducted using ExTaq (Takara Bio, Kusatsu, Japan) and amplicons were visualized on a 1.4% agarose gel.

### Reanalysis of Illumina WGS data

Publicly available Illumina WGS reads for non-stringy ‘Midas’, the Middle American stringy genotype ‘VAX3’, and the Andean stringy accession G4627 were downloaded and reanalyzed to assess if the duplication/insertion events identified for ‘Hystyle’ were shared by other non-stringy genotypes. The reference genome (G19833 v. 2.1; Phytozome 13: https://phytozome-next.jgi.doe.gov/info/Pvulgaris_v2_1) was augmented including the retrotransposon sequence. Reads were mapped against this augmented reference genome using bowtie2 (Langmead & Salzberg, 2012). Alignments were sorted by reference coordinates using Picard (https://broadinstitute.github.io/picard/) and visualized with the Integrative Genomics Viewer (https://software.broadinstitute.org/software/igv/) to identify reads of the accession Midas which spanned the retrotransposon boundary near *PvIND*.

## Results

### Transcriptional characterization

RNA-seq successfully characterized pod transcriptomes of four varieties across three pod development stages (Supplemental Fig. S1, Supplemental Table S6). Differentially Expressed Genes (DEGs) between stringy and non-stringy accessions were considered to identify gene ontology (GO) enrichments for pod string formation (Fig. 2, Supplemental Fig S1). A total of 237 genes were obtained from the R8 stage (Pod fill; Fernández et al. 1983) vs R7 stage (Initial pod development) comparison, 409 DEGs from the R9 (Pod maturation) vs R8 comparison (Pod fill), and 230 DEGs for the R9 vs R7 comparison. Functional enrichment analysis of GO terms of stringy accessions (Supplemental Fig. S2) showed activity of phosphatidylinositol phosphate kinases, N-acetyltransferases, phosphotransferases and union of transition metal ions as the most representative between stages R8 and R7. GO terms related to transferase, phosphotransferase and carbohydrate activity were mainly registered in the comparison R9 vs R8 in both gene sets. The enriched categories for R9 vs R7 included tetrapyrrole binding, oxidation-reduction activity, actin binding, binding to rRNA and nucleotidyl-transferase activity. Among all stringy accessions, membrane coat categories are included as enriched between R8 and R7, while the R9 vs R8 comparison did not present changes in these categories. Photosynthetic components, thylakoids, and membrane proteins were enriched in the comparison of stages R9 and R7 between stringy and stringless accessions.

**Fig 2.**
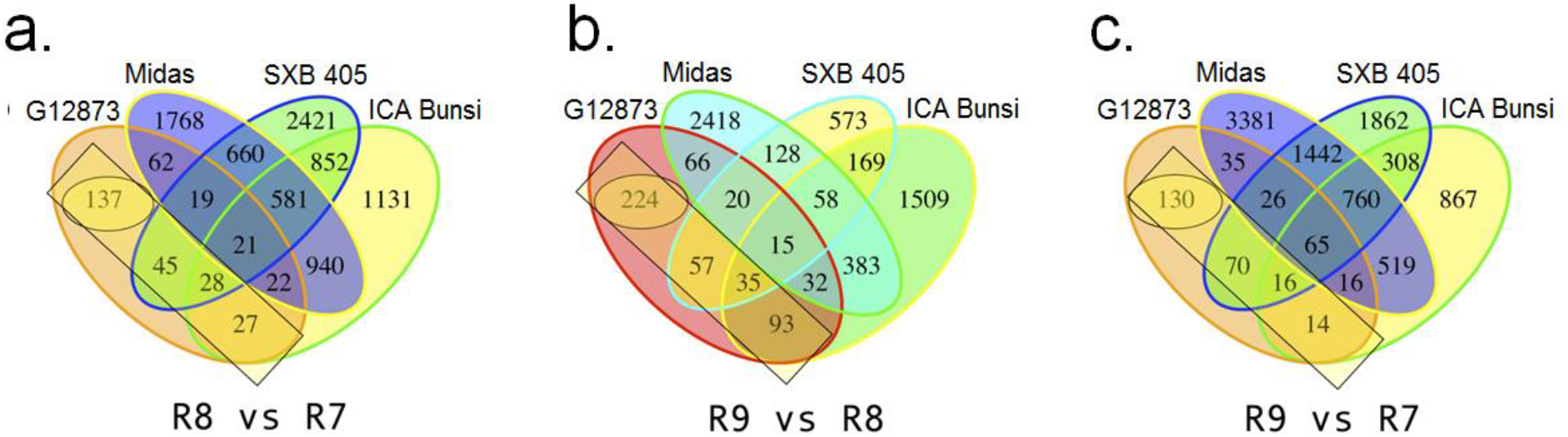
Venn diagram of differentially expressed genes (DEGs) observed for G12873, ICA Bunsi, SXB 405 and Midas when comparing the transcriptome at the indicated stages. The rectangles show DEG observed in G12873 and ICA Bunsi or SXB 405 or both, but not Midas.

Among the four Pv02 candidate genes, *PvIND* (Phvul.002G271000.1, Supplemental Figs. S3,S4) was statistically significantly more strongly expressed in the stringless accession than in the three stringy accessions (*p*=6.8*10^−6^, ANOVA of linear model). The results were independently significant at each maturity stage (R7: *p*=1.0*10^−4^; R8: *p*=4.3*10^−3^; R9: *p*=1.2*10^−5^; *t*-test). *PvIND* expression was non-overlapping between stringy and stringless varieties within each maturity stage. Of the other candidate genes, the NAC transcription factor Phvul.002G273100.1 was differentially expressed at the latest maturity stage (R9), with 3.6-fold higher expression in Midas than in the stringy varieties on average (*p*-value=0.01, *t*-test). Phvul.002G273100.1 expression at the other maturity stages individually and cumulatively across all stages was insignificantly different (*p* > 0.05, *t*-test). Similarly, the two other Pv02 candidate genes were insignificantly differentially expressed between phenotypic categories at any maturity stage.

### *PvIND* expression is predictive of string formation

RT-qPCR of non-stringy/stringy revertant pairs determined that *PvIND* expression was highly significantly correlated with pod strings across both sampling time points (Fig. 3). At five days after flowering, *PvIND* was approximately five-fold more expressed in stringless accessions than in stringy revertants, whether the control gene was *Act11* (4.9-fold difference, *p*=8.5*10^−5^) or *Ukn1* (5.3-fold difference, *p*=4.5*10^−4^). By 21 days after flowering, *PvIND* expression was 48-fold higher in stringless types when *Act11* was used as a reference (*p*=2.5*10^−6^) and 41-fold greater when *Ukn1* was used (*p*=1.3*10^−6^). In contrast, the three candidate genes of the NAC and MYB families were not significantly differentially expressed between phenotypic classes at either time point, regardless of which reference gene was used as a control. The minimum *p*-value of any of these comparisons was 0.06 (‘*NAC 2’* vs. *Ukn1*, 21 DAF).

**Fig 3.**
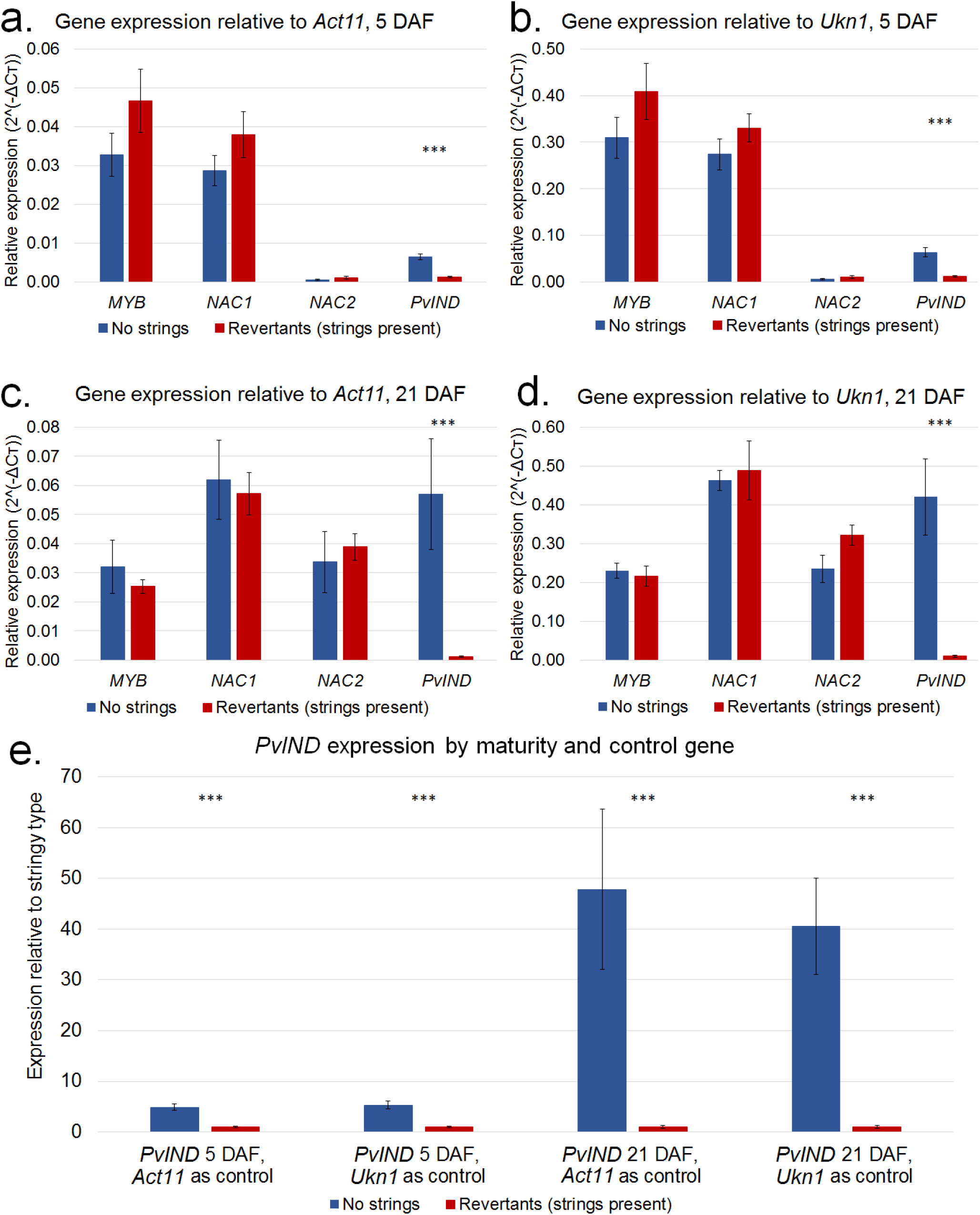
Significant differences in *PvIND* expression exist between cultivars with and without pod strings. At five days after flowering (DAF), a highly significant difference exists in *PvIND* expression whether a) *Act11* or b) *Ukn1* is used as a reference. No significant expression difference exists for other candidate genes. c)-d) At 21 DAF, the difference in *PvIND* expression has increased relative to each reference gene. No significant difference exists in expression of other genes. e) *PvIND* expression from previous panels re-scaled, relative to expression in stringy revertants. The difference in *PvIND* expression is greater later in pod maturity, and shows similar patterns regardless of reference gene. Three asterisks indicate *p*<0.001, no asterisks indicate *p*>0.05. Error bars represent standard error of the mean of six replicates of each phenotypic class. *PvIND*: Phvul.002G271000; *NAC 1*: Phvul.002G271700; *NAC 2*: Phvul.002G273100; *MYB*: Phvul.002G269900, *Act11*: Phvul.008G011000; *Ukn1*, Phvul.011G023200.

### *PvIND* expression associated with changes in cell identity

Major differences in secondary cell wall development distinguished stringless and stringy revertant types (Figs. 4,5). Revertant vascular sheaths were nearly identical to those of wild or dry beans (Fig. 4b, Prakken 1934, Parker 2020). In revertant pods, the vascular sheath is primarily composed of 3-6 layers of thickly-lignified fiber cells. At the center of the sheath is a narrow strip of weakly lignified cells approximately two cells wide, known as the dehiscence zone. In contrast, stringless varieties produce dehiscence zone-like cell layers throughout the vascular sheath. These are lignified but lack secondary cell wall thickening and have a small fraction of the cross sectional cell wall surface area of fiber cells. The total area of the vascular sheath is also reduced, with only 1-3 cell layers in non-stringy varieties. Only one type, the Prevail revertant, showed strong pod wall fiber along with pod strings. The stringless form of Prevail also shows subtle wall fiber deposition (Fig. 5).

**Fig 4.**
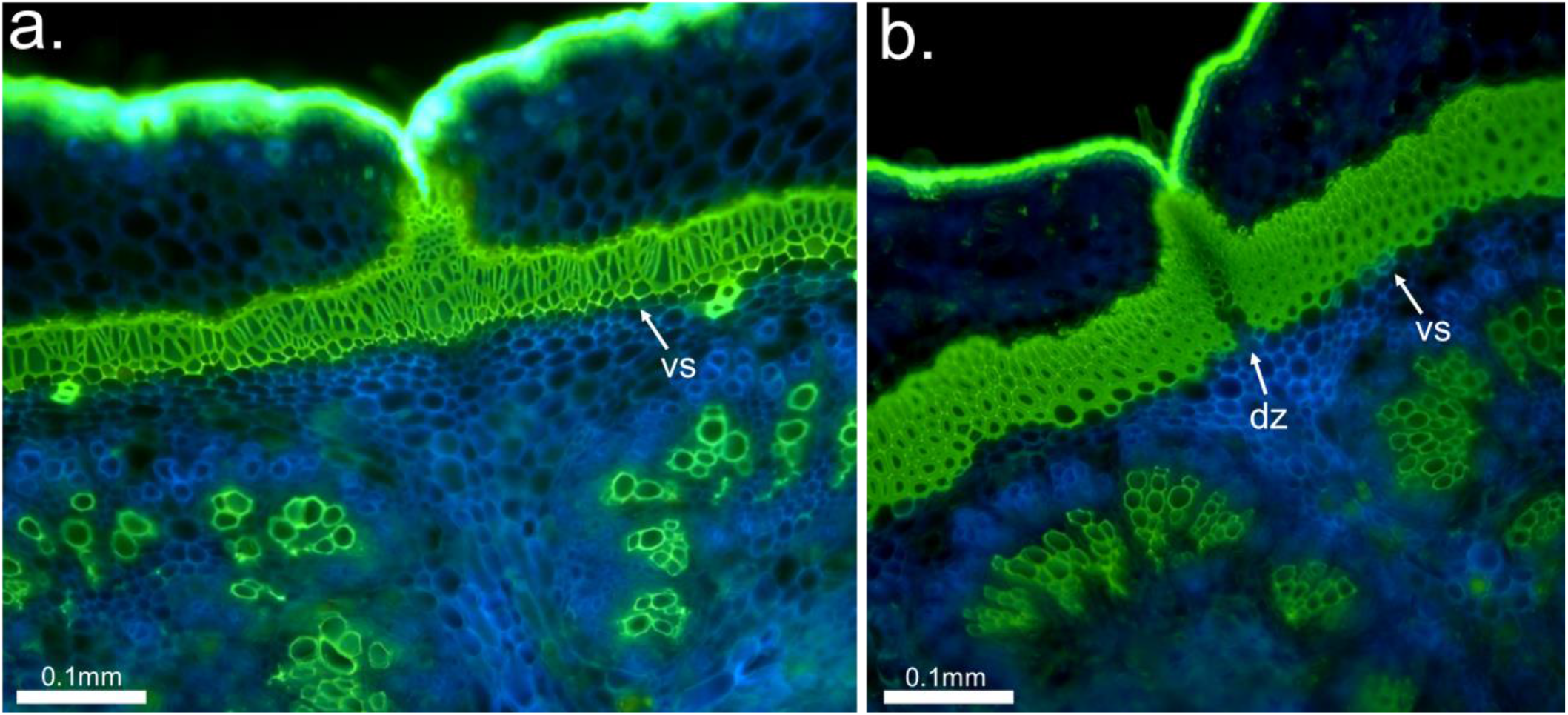
Anatomical comparison of a) *PvIND*-overexpressing (non-stringy) and b) revertant (stringy) pods. In a) non-stringy accessions, vascular sheaths (vs) show little to no secondary cell wall development and have just 1-3 lignified cell layers, whereas b) stringy revertants have strong secondary thickening and 3-6 fiber cell layers, except in the dehiscence zone (dz) along which dehiscence occurs in susceptible varieties. In non-stringy types with *PvIND* overexpression, over-specification of weak dz-like cells throughout the vs leads to the lack of pod suture string. Samples of a) Nampa-20 ‘BBL156’ and b) Nampa-3 ‘Hystyle’ shown.

**Fig 5.**
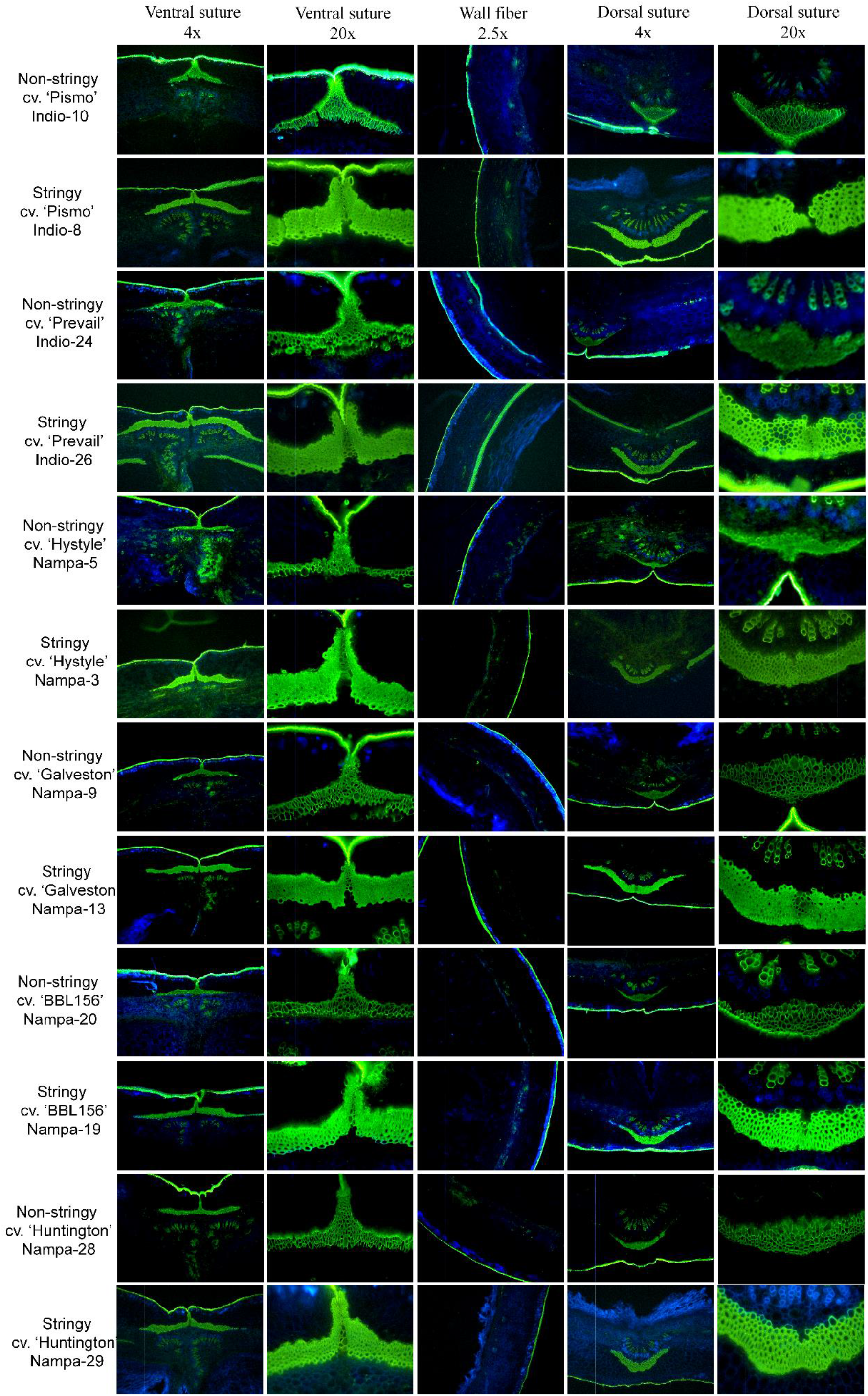
Anatomy of stringless and revertant-stringy pairs. In all cases, the non-stringy type shows a near total absence of secondary cell wall thickening. All revertants show this weakly lignified cell type only in a narrow band of cells in the dehiscence zone, while the majority of cells have strong secondary thickening, as well as a somewhat greater number of cell layers in the vascular sheath. In five of six revertants, there is no notable effect on fiber deposition in the pod walls. Reversion in Prevail, however, was associated with a major increase in pod wall fiber deposition. This indicates that *PvIND* may, in some cases, be a regulator of pod wall fiber development.

### Conservation in *PvIND* homolog promoters

Three main regions of high similarity were identified among promoters of *PvIND* homologs. These spanned from approximately 1,514- 1,643 bp, 906-1,048 bp, and 293-430 bp before the *PvIND* transcription start site, and were enumerated as motifs 1, 2, and 3, respectively (Fig. 6). While motifs 1 and 3 were conserved among all legumes analyzed, motif 2 was conserved only among the Phaseoleae. At the middle of these were core sequences with very high conservation. In motif 1, the sequence CCCTAGGATTTCAGTGC was identified without substitution or gaps for 17 of 21 gene models, while in the other four there were no more than two SNPs. Motif 3 included the sequence (ATGCTTTTTGCAGTSASW(C)_0-1_CCCCTTTCAGTAAAAAC) across all species with above-ground pods. The end of this conserved sequence and 50bp immediately following it were predicted to have enhancer activity by EnhancerPred, while this was not the case anywhere else in the 2.5kb region upstream of *PvIND*. The comparison of *IND* homologs in common bean and Arabidopsis indicated that *PvIND* most closely clusters with *AtIND* rather than other similar proteins (Supplemental Fig. S4).

**Fig 6.**
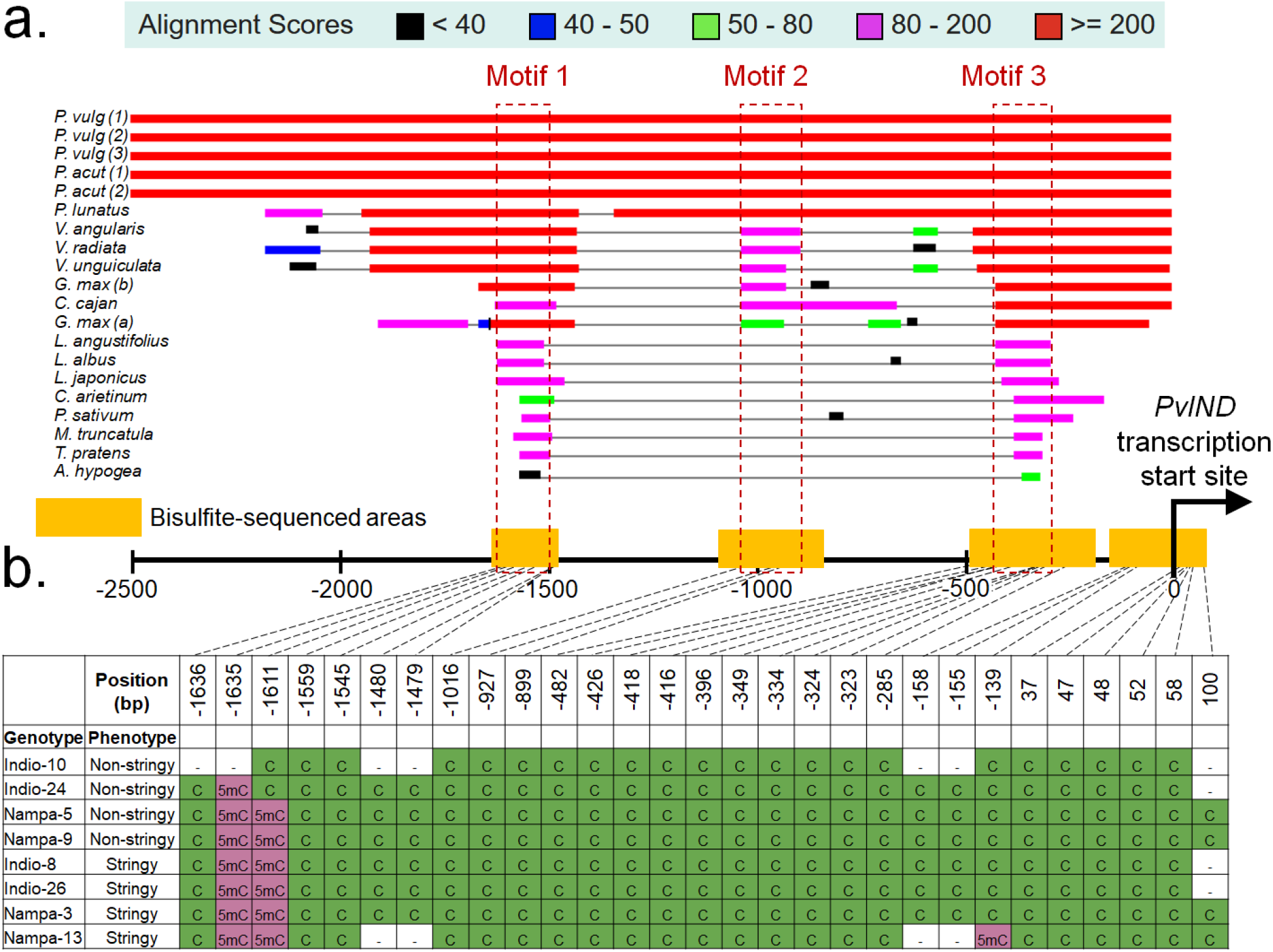
Bisulfite sequencing of conserved PvIND promoter elements. a) Multiple alignment of the 2.5kb sequence upstream of legume *PvIND* orthologs revealed three regions of conserved non-coding sequence across the legume family. Motif 3 was predicted by EnhancerPred (Jia and He 2016) to have transcriptional regulatory activity. Extreme conservation of these motifs indicated that they may regulate *PvIND* transcription. Since no DNA sequence variations in these regions differentiate phenotypic categories, b) bisulfite sequencing was conducted on non-stringy/stringy revertant pairs. Methylation status of all sequenced CG and CHG positions in promoter motifs are displayed, with green representing unmethylated sites and purple showing methylated sites. No fixed differences in methylation were identified between stringy and non-stringy types in these regions, indicating that other factors affect *PvIND* expression.

### DNA methylation of *PvIND* promoter is not predictive of pod strings

Bisulfite sequencing returned reads at three conserved motifs upstream of *PvIND*, 246 bp flanking the transcriptional start site, 270 bp in the gene body, and 48 bp of 3’ UTR. Relative to the transcription start site, these ranged from −1,652 to −1,474 bp, −1,136 to −906 bp, −489 to −215 bp, −186 to 102 bp, 406 to 676 bp, and 1,187 to 1,234 bp. No surveyed positions were predictive of pod phenotype (Fig. 6b). Methylated cytosines existed in motif 1, with two methylated residues; as well as the gene body, with two identical methylated cytosines in both stringy and non-stringy variants of Hystyle. Across surveyed cytosines, methylation patterns tended to be highly consistent among accessions, and at no position did methylation status predict pod string phenotype.

### *PvIND* duplication and retrotransposon insertion

HiFi/BioNano mapping identified several unique features of the non-stringy accession ‘Hystyle’ in the region surrounding *PvIND*, including tandem duplication of *PvIND* and insertion of a retrotransposon between the two copies (Fig. 7). PacBio HiFi sequencing averaged a depth of 45x, with median read lengths of 17.3 kb. These sequences were combined with the BioNano mapping results to create a single scaffold containing the full 500 kb region of Hystyle between the *PvIND* flanking markers identified by Hagerty et al. (2016). All seven available *Phaseolus* reference genomes are of accessions with stringy pods, with a single copy of *PvIND* and no retrotransposon in the surrounding region. In contrast, Hystyle contains a 12kb tandem duplication of *PvIND* and its upstream promoter. The duplicated copies (*PvINDa* and *PvINDb*) are identical in putative transcribed sequence. Further, the second tandem repeat included a *Ty1-copia* family retrotransposon insertion approximately 10 kb upstream of *PvINDb* and 400 bp downstream of *PvINDa*. The retrotransposon is 2,760 bp in length, and includes insertion site repeats and terminal inverted repeats typical of the transposable element family. Finally, CT repeats of 278 bp and 170 bp immediately follow *PvINDa* and *PvINDb*, respectively, whereas these repeats are no longer than 140 bp in any other *Phaseolus* reference genome. Reanalysis of previously sequenced Illumina data for the non-stringy accession ‘Midas’ (Lobaton et al., 2018) also supports the retrotransposon insertion for this accession (Supplemental Fig. S5).

**Fig 7.**
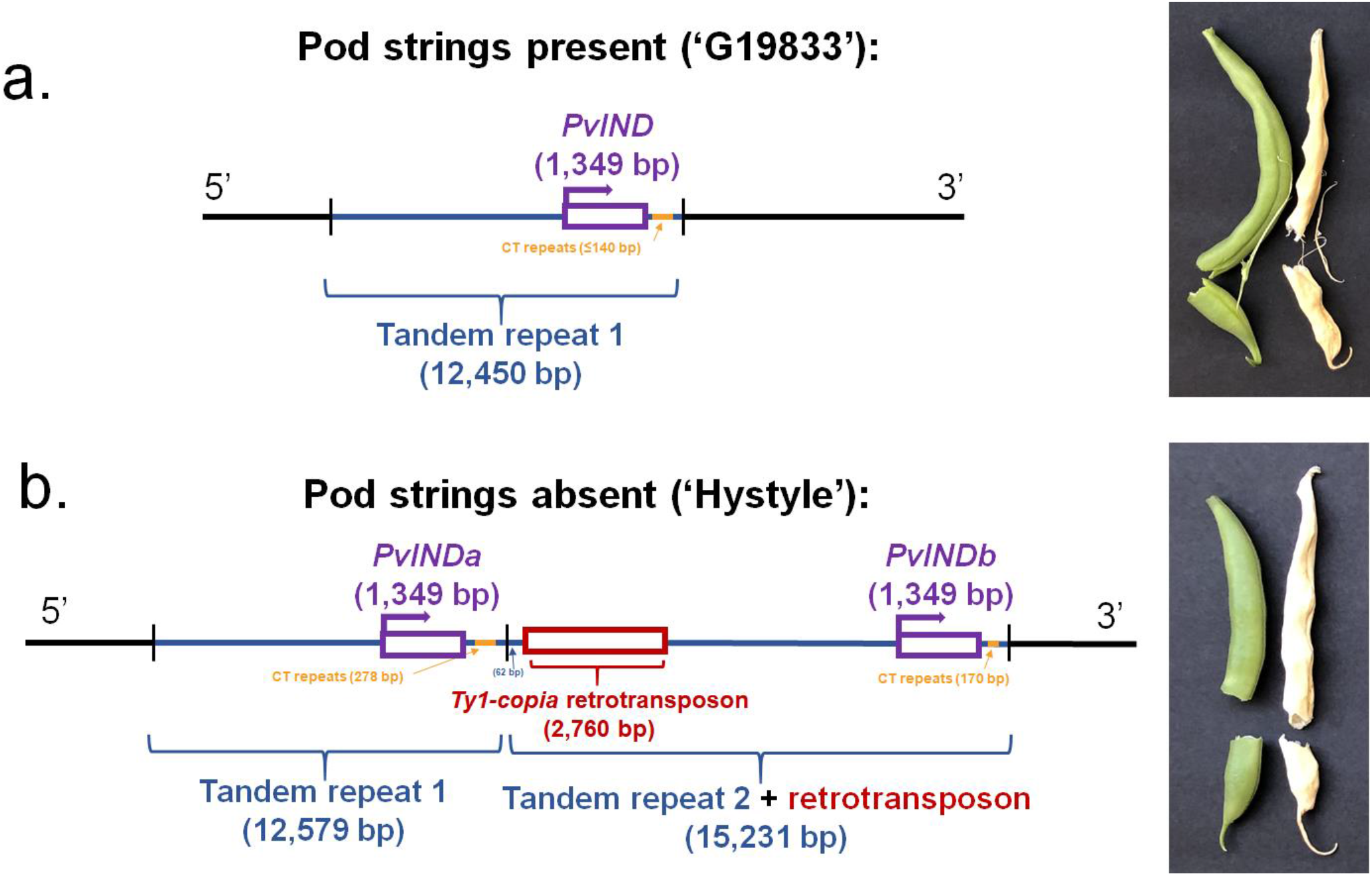
*PvIND* sequence structure between stringy and non-stringy accessions. a) Stringy accessions such as G19833 have a single copy of *PvIND* without retrotransposon. In contrast, b) non-stringy accessions such as Hystyle show a tandem duplication of *PvIND*, with a retrotransposon inserted at the beginning of the duplicated region. These sequence features may cause the *PvIND* overexpression associated with loss of pod strings.

### Sequence variation associated with pod strings in revertants and other accessions

Among 100 diverse accessions, tandem duplication of *PvIND* was always associated with retrotransposon insertion and vice versa. Tandem duplication and retrotransposon insertion were predictive of pod string phenotype, without phenotypic overlap between genotypic categories (Fig. 8, Supplemental Fig. S6, Supplemental Table S7). This determined that previously reported “recombination” between *PvIND* and pod strings (Gioia et al. 2013) may have been based on errors in phenotyping due to assumption of co-inheritance of pod and wall fiber (Koinange et al. 1996), as our results clearly matched genotype and phenotype in re-analyzed RILs (Supplemental Table S7). Intriguingly, stringy revertants did not include the tandem duplication and retrotransposon insertion, unlike their isogenic non-stringy lines, indicating that the reversion process involves loss of these sequence features.

**Fig 8.**
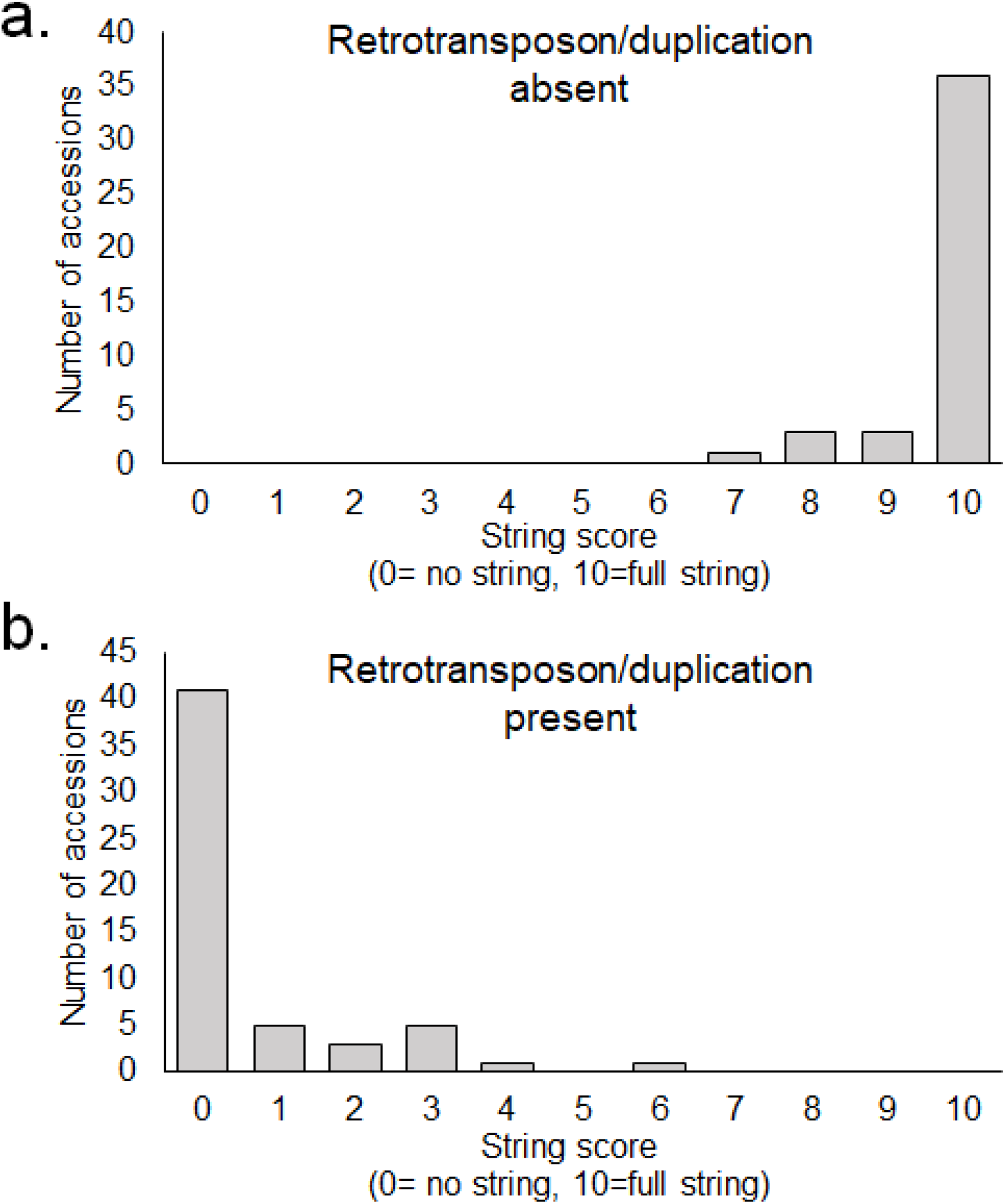
Among 100 diverse accessions of common bean, reduced pod string score is strongly related to the *Ty1-copia* retrotransposon presence near *PvIND* and duplication status of the gene. a) accessions lacking these sequence features tend to produce strong suture strings, while b) accessions with the retrotransposon insertion and gene duplication produce little to no pod strings.

## Discussion

Our results demonstrate that the loss of pod strings in common bean is associated with duplication of *PvIND* and a retrotransposon insertion between the two gene copies. Accessions with these sequence features consistently express *PvIND* transcripts at abundances at least 40- fold higher than isogenic revertant lines, which lack these features. *PvIND* expression level between stringless and revertant lines is uniformly related to over-specification of weak pod dehiscence-zone tissue throughout the vascular sheath, in cell types that would otherwise produce strong secondary cell wall deposition.

During domestication and breeding of autogamous species, the vast majority of new character states are mediated by recessive alleles based on loss-of-function mutations. This is the case in common bean, for key domestication traits such as determinacy, photoperiod insensitivity, loss of seed dormancy, white-seededness, and reduction in pod shattering (Parker and Gepts 2021). In contrast, current results show that a major gain in *PvIND* expression has occurred in the development of stringless forms, paralleling the dominant nature of the apomorphic allele. Our RNA-seq results show that Midas has by far the most unique pattern of gene expression in pods among the four analyzed accessions. This is consistent with the idea that loss of pod strings led to more extensive pod remodeling than the initial loss of pod shattering, a core element of legume domestication (Parker et al. 2021a). Our RNA-seq results parallel those of Di Vittori et al. (2021), who also identified *PvIND* among a list of genes differentially expressed between Midas and G12873. The distant relationship between these accessions, which descend from distinct gene pools (Andean vs. Middle American, respectively; Parker and Gepts 2021), necessitated an analysis of gene expression in a controlled genetic background. Our RT-qPCR results conducted in six isogenic backgrounds demonstrate unequivocally that differential *PvIND* expression is not only correlated with string deposition but is also qualitatively predictive of string formation among all samples and at both sampled time points. This is compelling evidence that *PvIND* expression regulates pod string formation in common bean.

We next analyzed the anatomical effects of *St* reversion in the isogenic stringless/revertant pairs. In all cases, stringless accessions demonstrated an expansion of the weak cells of the dehiscence zone throughout the vascular sheath. In Arabidopsis, *IND* regulates the development of the valve margin layer where breaking occurs (Girin et al. 2010; Liljegren et al. 2004), suggesting that *PvIND* specifies the comparable weak dehiscence layer in common bean. Loss of pod strings could therefore be the result of ectopic *PvIND* overexpression throughout the bundle sheath. Methods such as RNA *in situ* hybridization and laser capture microdissection RT-qPCR could be useful to test this possibility. Additionally, five of six revertants showed no appreciable increase in pod wall fiber deposition, indicating that the effect of *St* is typically specific to the string region in most genetic backgrounds. The identification of wall fiber deposition in the ‘Prevail’ revertant indicates that *PvIND* may prevent pod wall fiber development in some environments and/or genetic backgrounds.

Several regions upstream of *PvIND* have been conserved base-by-base over tens of millions of years of legume evolution. This indicates that they play a role critical to the survival of many wild legumes. Bisulfite sequencing of these and other regions in and around *PvIND* found no patterns of DNA methylation predictive of pod string formation, ruling out their role in the reversion and regulation of string formation.

The *PvIND* tandem duplication and retrotransposon insertion are both logical drivers of *PvIND* overexpression in non-stringy types. Gene duplication frequently leads to enhanced expression due to increased gene copy number (Gemayel et al. 2012), while transposable elements typically contain transcription-enhancing motifs with long-range effects (Lisch 2013, Hirsch and Springer 2017). Transposable elements frequently lead to ectopic gene expression, which could explain the gain of dehiscence zone cell identity throughout bundle sheath layers in stringless varieties. The existence of the retrotransposon in only the second tandem repeat indicates that its insertion occurred after gene duplication. Intermediate forms with *PvIND* duplication but without the retrotransposon would be useful to separate the role of each sequence feature, although these types have not yet been identified.

The uniform loss of the *PvIND*-associated retrotransposon across all eight stringy revertant lines is an interesting find. While excision of DNA transposons occurs regularly, this process is not known to occur among retrotransposons, which instead replicate via reverse transcription. Unequal crossing over between the tandem duplications is one possible mechanism that could explain this loss, but further sequencing and characterization of the region will be important to better understand the genetic basis of the reversion process.

## Data availability

The 500kb Hystyle sequence between the *PvIND* flanking markers will be submitted to NCBI. FASTA of bisulfite-converted DNA will be submitted to DataDryad.

## Author contributions

Conceptualization and interpretation: PG, AHE, TP, EA, JRM; Molecular biology: TP, JC, LLS, SK, SL, SN, TOF; Bioinformatics: VL, JD, TP, JC, AHE; Microscopy: TP, SL, TOF, JJ; Funding acquisition: PG, AHE, TP; Writing – Original Draft: TP, JC, PG. All authors revised and approved the final manuscript.

## Acknowledgements

This work was supported by funding from Clif Bar/Seed Matters, Lundberg Family Farms, Kirkhouse Trust, Henry Jastro Graduate Research Scholarships, the Universidad de los Andes FAPA initiative led by the Vice-presidency of Research and Knowledge Creation, CAPES Postdoctoral Program/Process n*88881.170593/2018-01, UC MEXUS grant CN-17-46, and USDA-NIFA Regional Hatch project W3150. Dan Wahlquist and Syngenta generously allowed us to collect stringless/revertant types at their facilities.

## Supplemental Information

### Supplemental Figures

**Supplemental Fig. S1.**
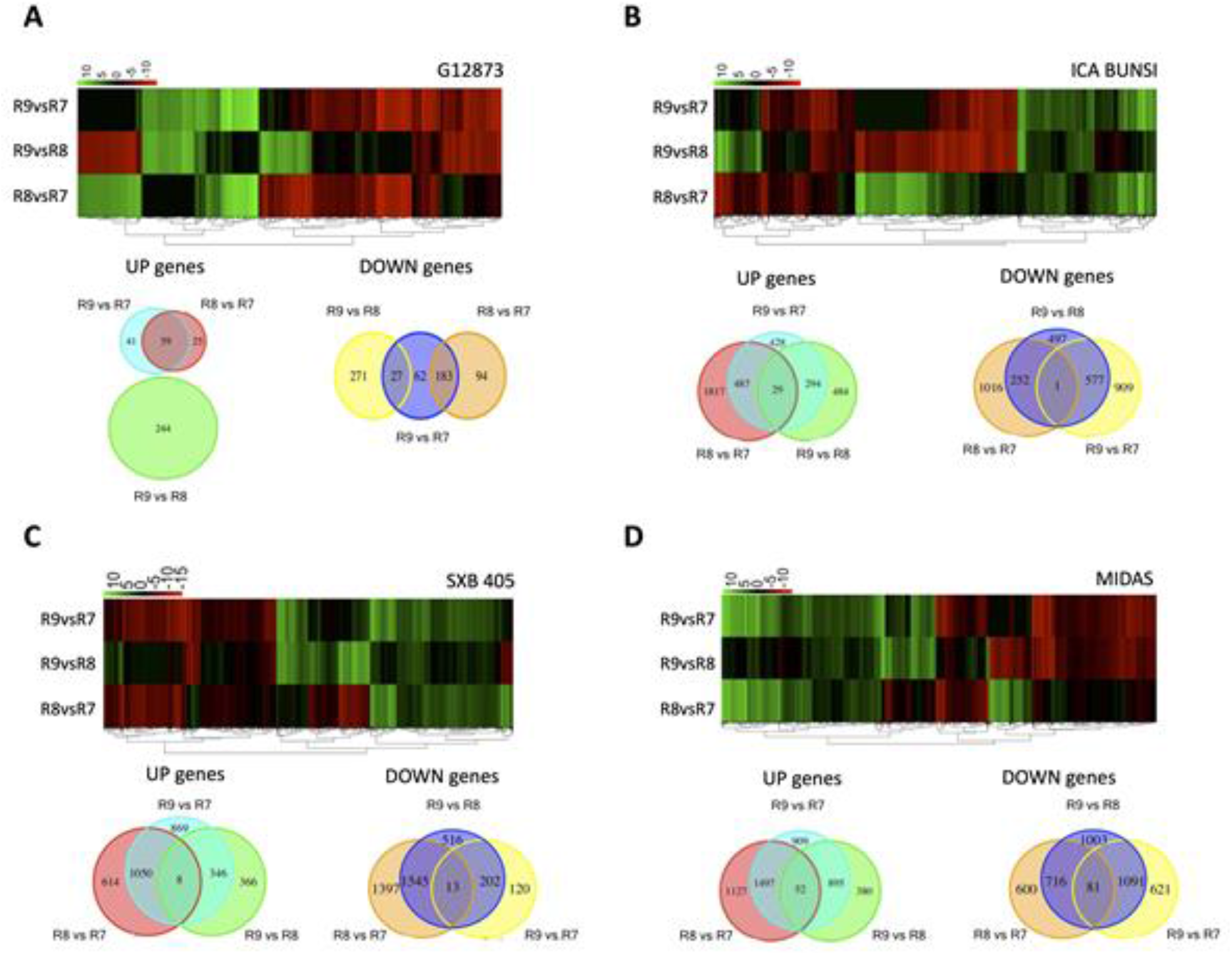
Transcriptome profile of pod development from a) wild dehiscent *Phaseolus vulgaris* G12873, b) domesticated dehiscent cultivar ICA Bunsi, c) domesticated indehiscent SXB 405, and d) domesticated stringless Midas. log2FC of differentially expressed genes from the pairwise comparison of R8 vs R7, R9 vs R8 and R8 vs R7 are shown in the heatmaps and grouped in Venn diagrams selecting up or down regulated genes by developmental stage. Green bars represent up-regulated genes, red shows down-regulated genes and black bars display genes without expression changes.

**Supplemental Fig. S2.**
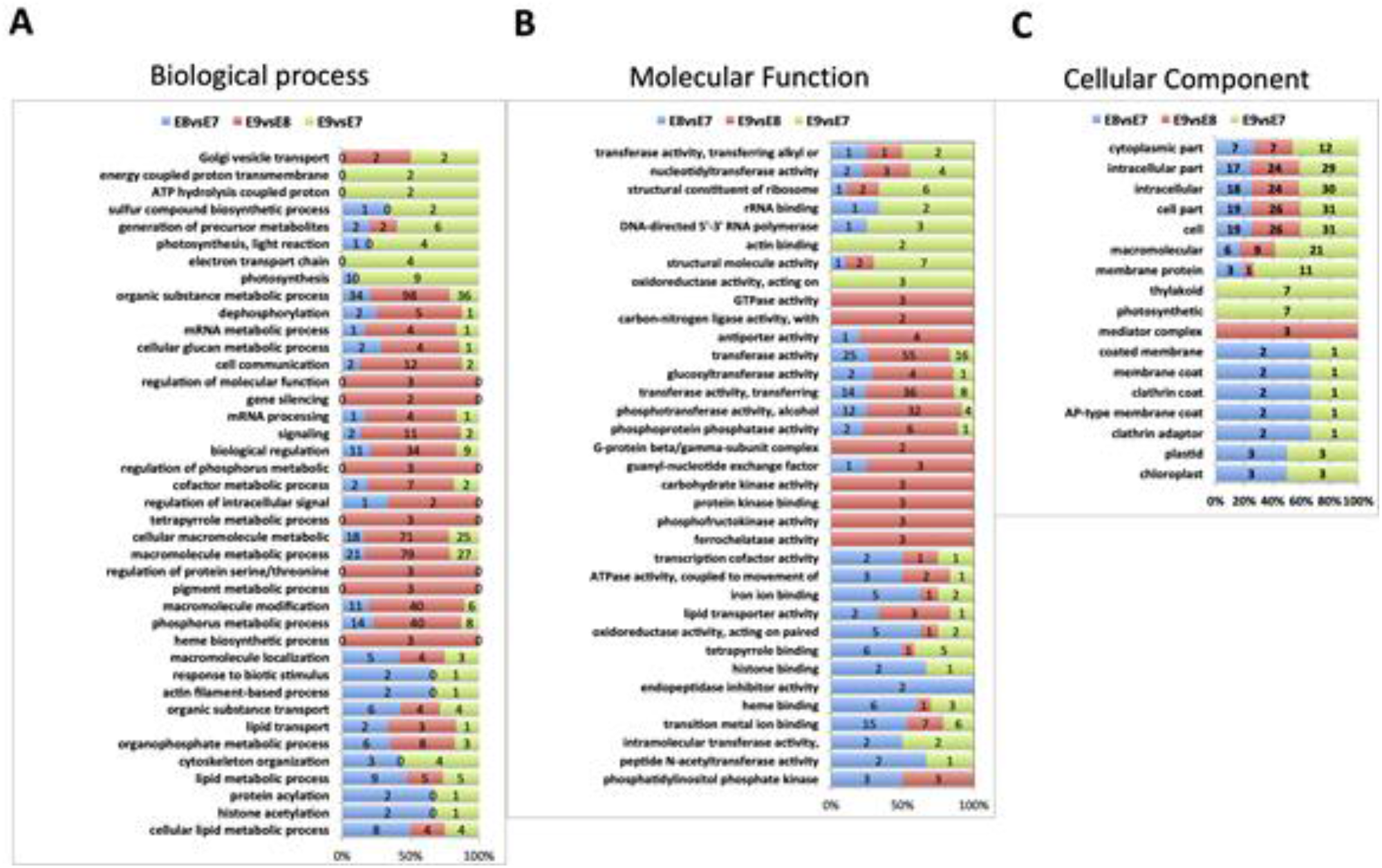
Gene ontology in terms of Biological process (A), Molecular Function (B) or Cellular Component (C) enriched from DEG in G12873, ICA Bunsi and SXB 405 but not in Midas.

**Supplemental Fig. S3.**
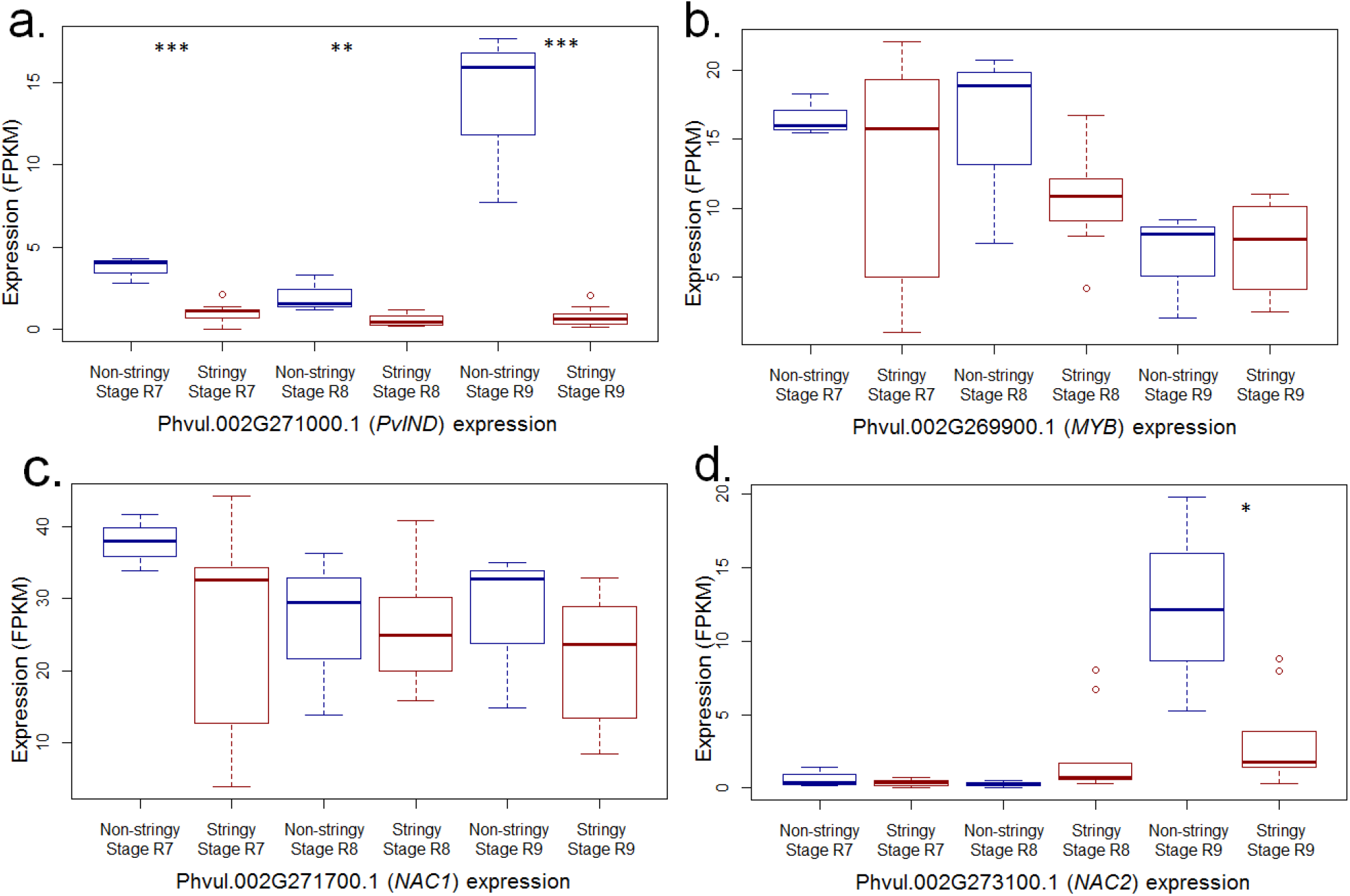
RNA-seq of four candidate genes between *St* flanking markers on Pv02. A) At all pod maturity stages, expression of Phvul.00G271000.1 (*PvIND*) in Midas (non-stringy) is significantly higher than in the three stringy varieties ICA Bunsi, SXB 405, and G12873 (*t*-test *p*<0.01). In contrast, expression of B) Phvul.00G269900.1 (*MYB*) and C) Phvul.00G271700.1 (NAC1) are insignificantly different between classes at all time points. D) Expression of Phvul.00G273100.1 (NAC2) is insignificantly different at the R7 and R8 sampling points, but is significantly more expressed in non-stringy Midas than in the stringy varieties at R9 (*t*-test *p*=0.01). Three asterisks indicate *p*<0.001, two asterisks indicate 0.001<*p*<0.01, one asterisk indicates 0.01<*p*<0.05, no asterisks indicate *p*>0.05

**Supplemental Fig. S4.**
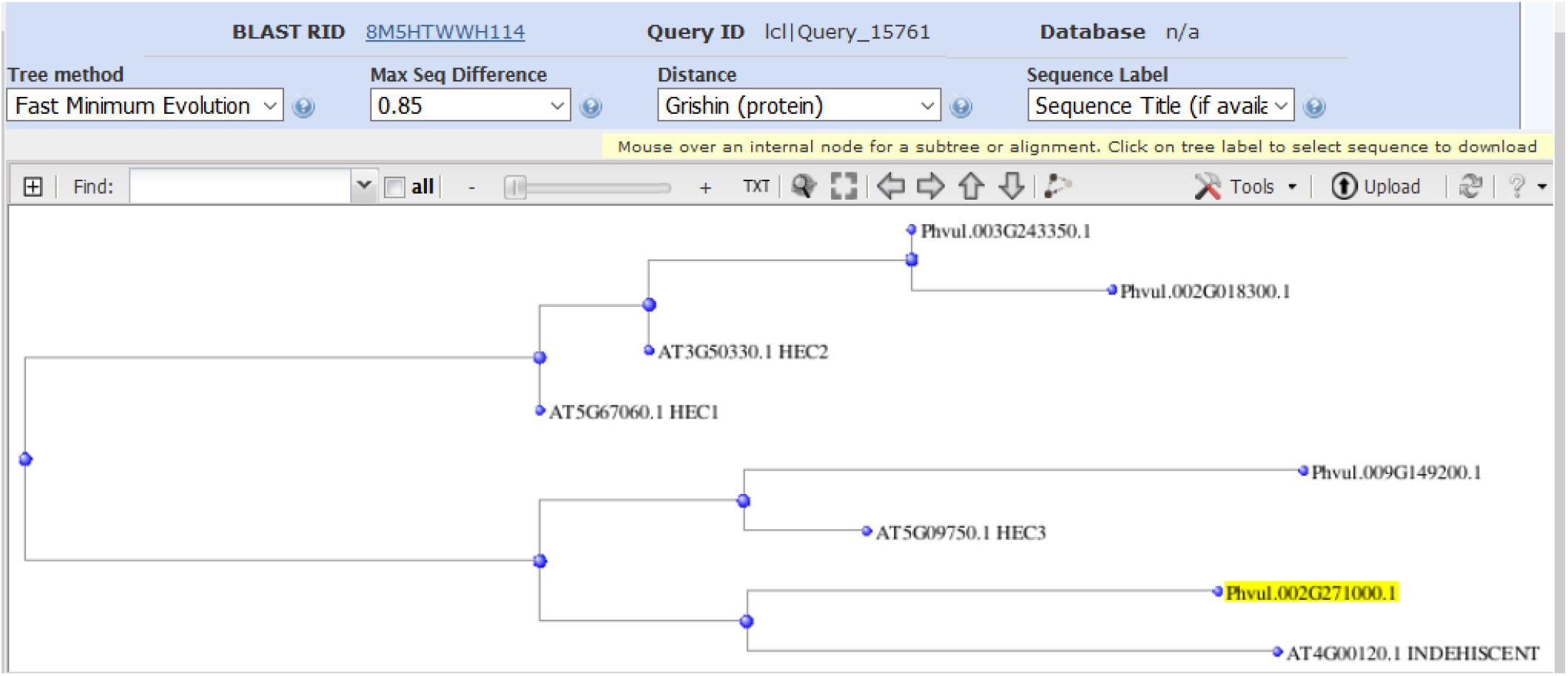
Distance tree of amino acid sequence data of Phvul.002G271000.1 (*PvIND*) and closest relatives in *P. vulgaris* and *A. thaliana*. Phvul.002G271000.1 clusters most closely with *INDEHISCENT* of Arabidopsis.

**Supplemental Fig. S5:**
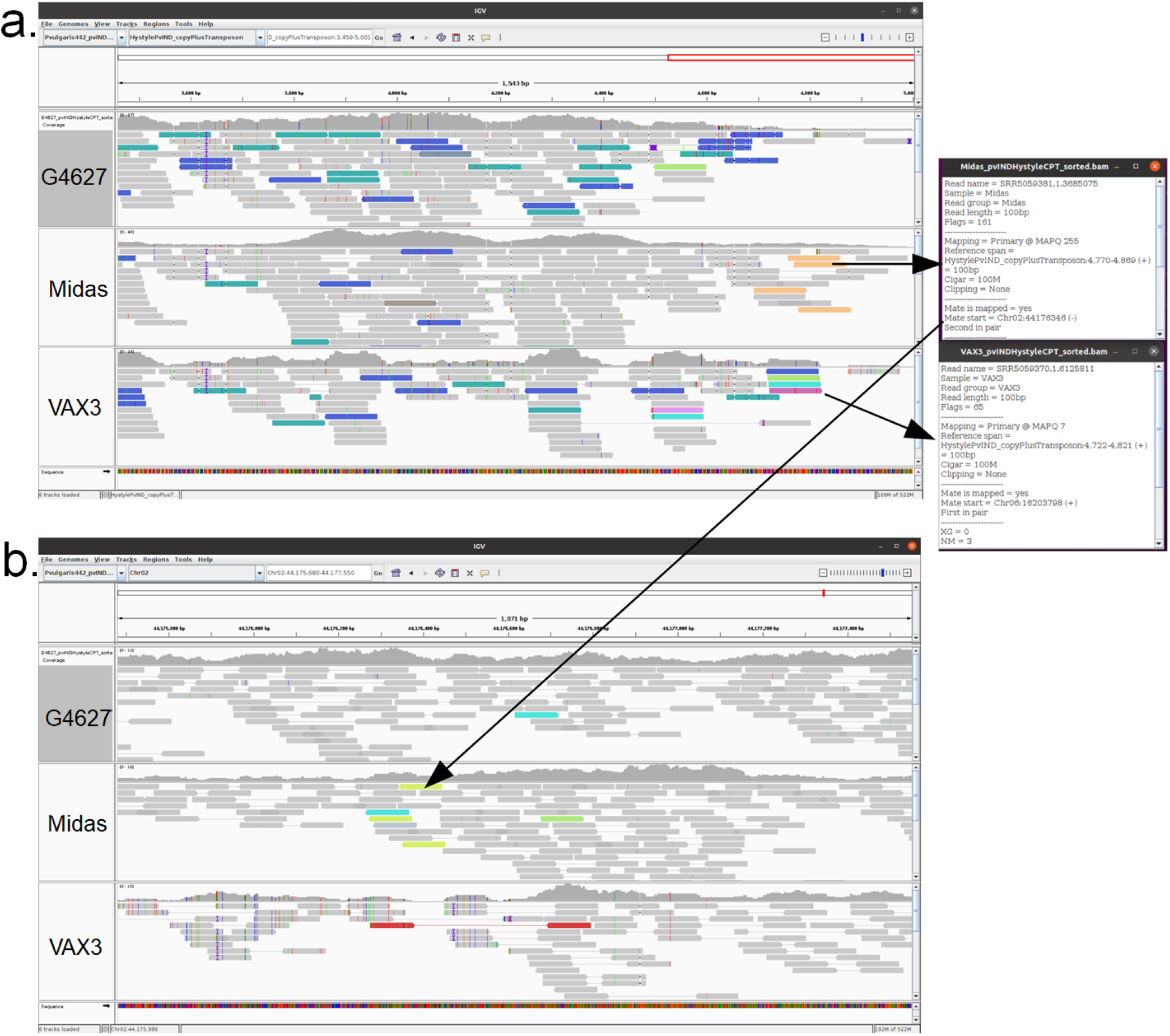
Reads previously sequenced from the genotypes G4627, Midas, and VAX 3, aligned to a) the *Ty1-copia* retrotransposon found near *PvIND* in the accession ‘Hystyle’, and b) to the sequence upstream of *PvIND* in the reference genome G19833, which lacks the retrotransposon. Colors other than gray indicate abnormal alignments. The right box shows an example of a Midas fragment for which one read maps to the end of the retrotransposon and the other end maps to the promoter of the reference copy of *PvIND* at 44,176,346. In contrast, G4627 has a depth close to zero at the end of the transposon, and for VAX 3 reads map to different chromosomes, suggesting a possible insertion of this transposon in a different location of the genome.

**Supplemental Fig. S6.**
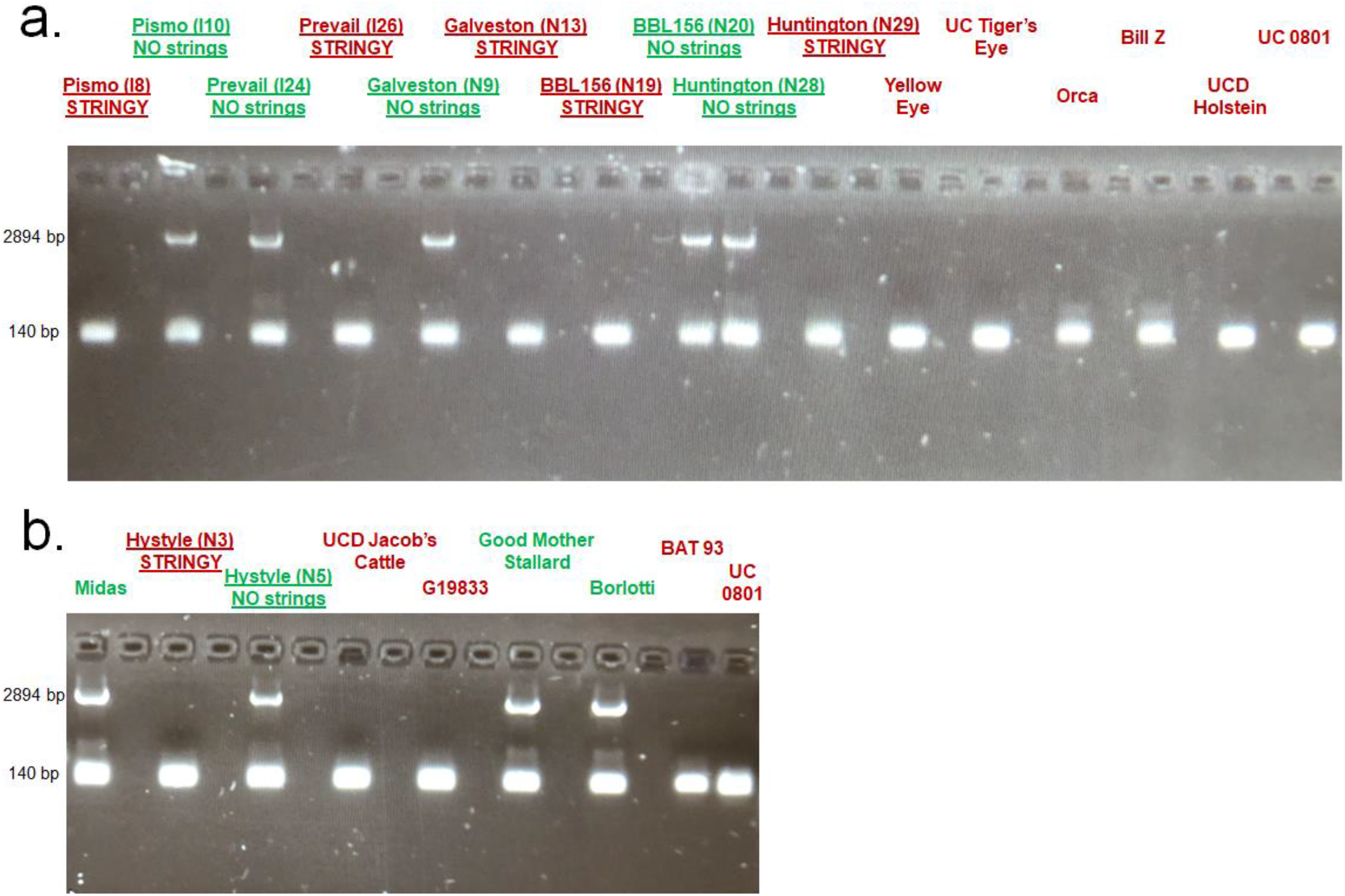
Amplification of retrotransposon-related locus near *PvIND*. The primer pairs BRT F1 + ART R5 span the retrotransposon site. In stringy accessions (labeled in red), the site contains no retrotransposon and only the 140 bp fragment is amplified. In non-stringy accessions (labeled in green) the retrotransposon is present, leading to a 2894 bp fragment from the second tandem repeat, while the first tandem repeat produces a 140 bp fragment. Isogenic stringless/revertant pairs are underlined.

### Supplemental Tables

**Supplemental Table S1.** Plant material information for the four *P. vulgaris* accessions used for RNA-seq.

| Accession | Gene pool | Wild / domesticated | Pod string | Shattering |
| --- | --- | --- | --- | --- |
| G12873 | Middle American | Wild | Strong string | Very high |
| ICA Bunsí | Middle American | Domesticated | Strong string | Intermediate |
| SXB 405 | Middle American | Domesticated | Strong string | Low |
| Midas | Andean | Domesticated | No string | None |

**Supplemental Table S2.** Pairs of stringy and non-stringy lines of the same genetic background.

| Sample code | Genotype | Phenotype |
| --- | --- | --- |
| Indio-8 | Pismo | Stringy |
| Indio-10 | Pismo | Not stringy |
| Indio-24 | Prevail | Not stringy |
| Indio-26 | Prevail | Stringy |
| Nampa-3 | Hystyle | Stringy |
| Nampa-5 | Hystyle | Not stringy |
| Nampa-9 | Galveston | Not stringy |
| Nampa-13 | Galveston | Stringy |
| Nampa-19 | BBL156 | Stringy |
| Nampa-20 | BBL156 | Not stringy |
| Nampa-28 | Huntington | Not stringy |
| Nampa-29 | Huntington | Stringy |

**Supplemental Table S3.**
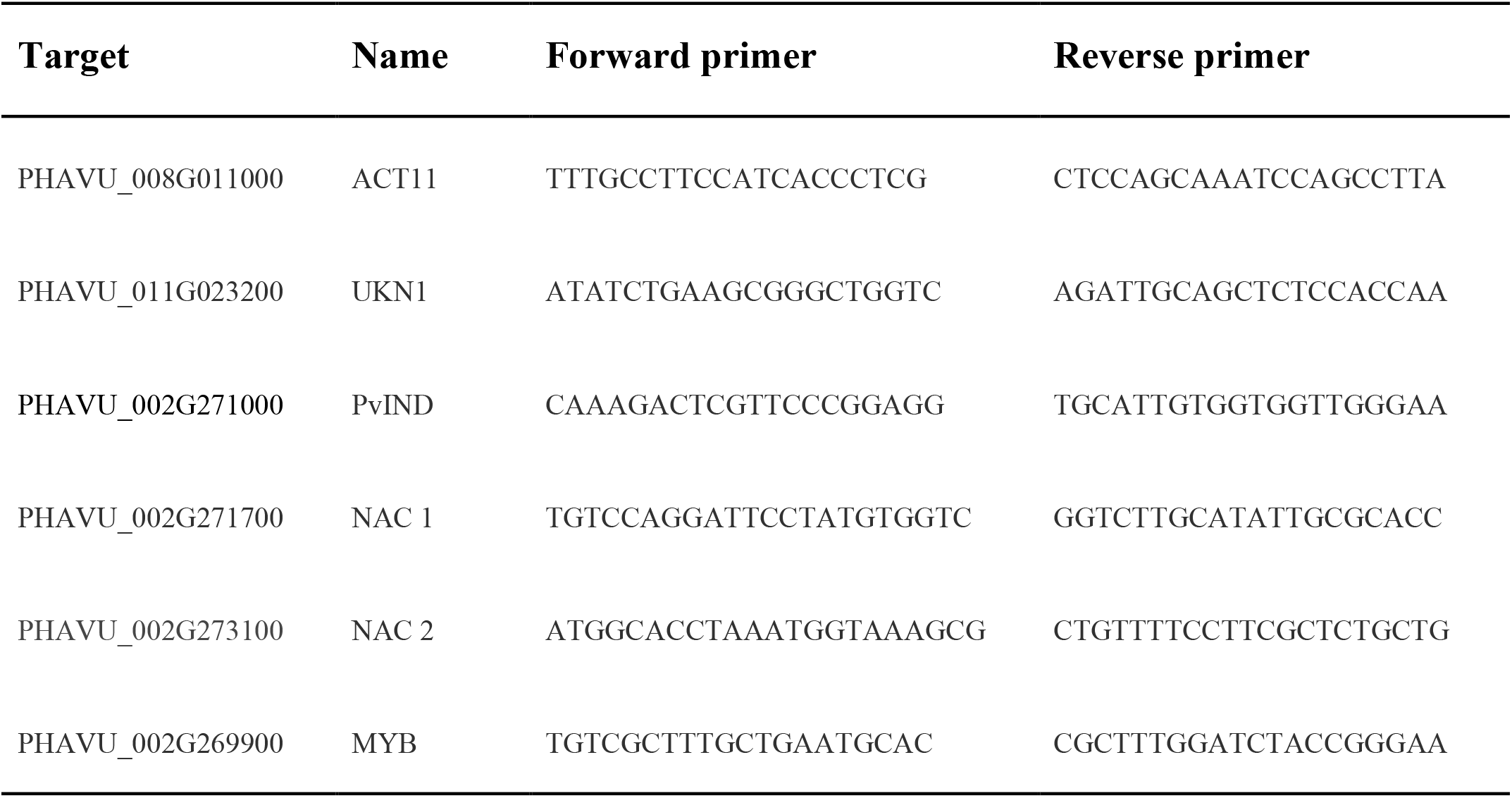
Primers used for RT-qPCR.

**Supplemental Table S4.**
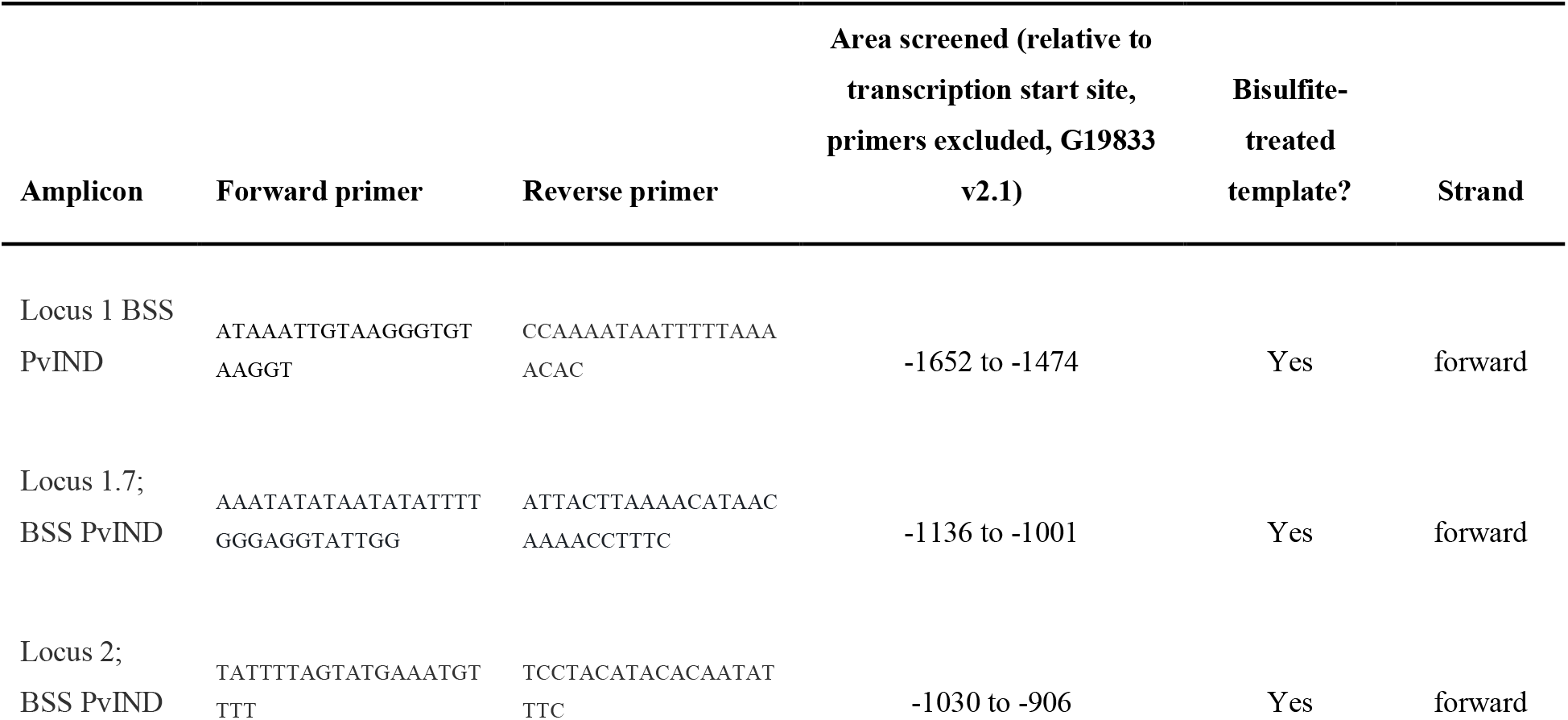
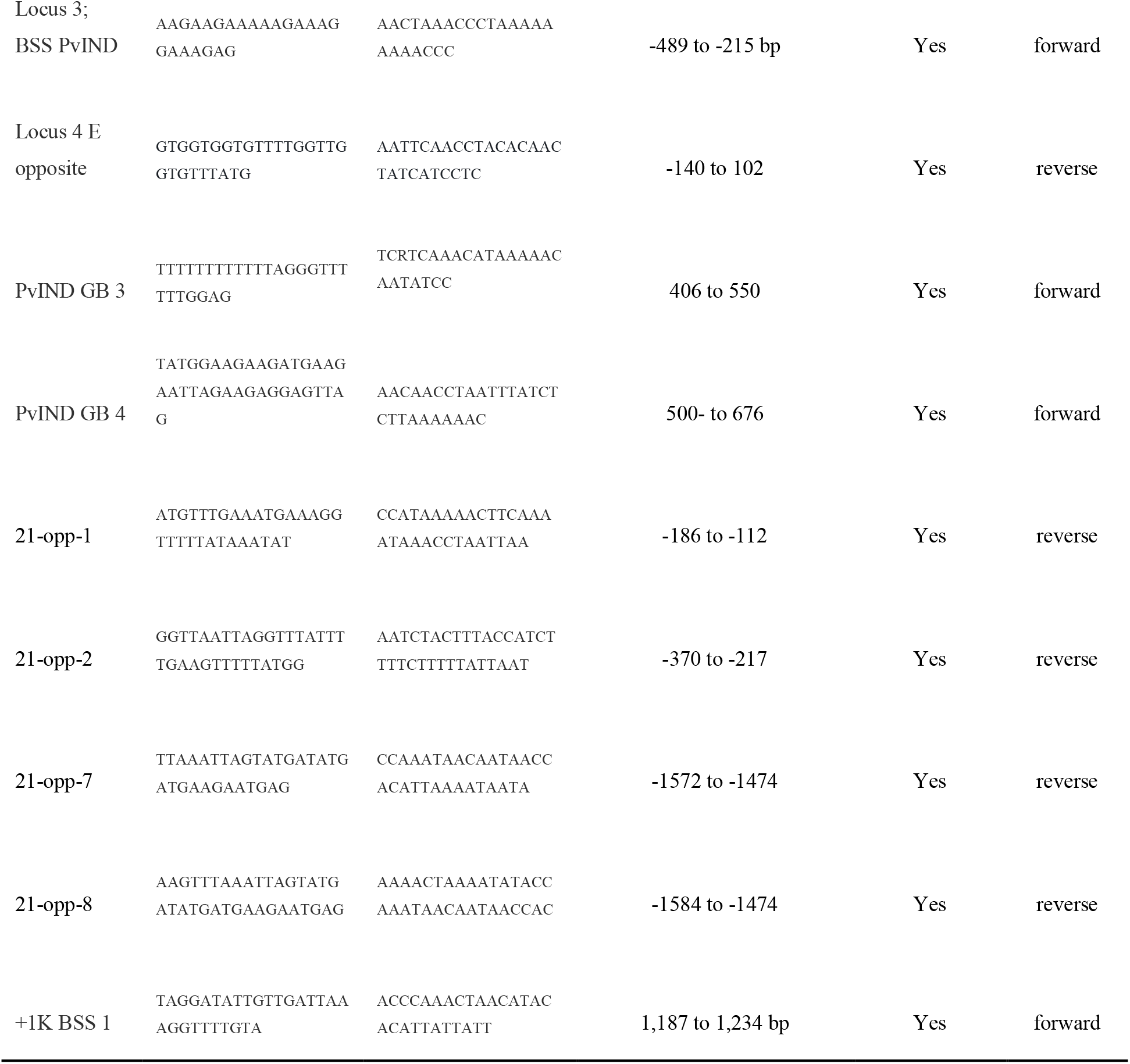
Summary of primers and amplicons used for bisulfite sequencing.

**Supplemental Table S5.**
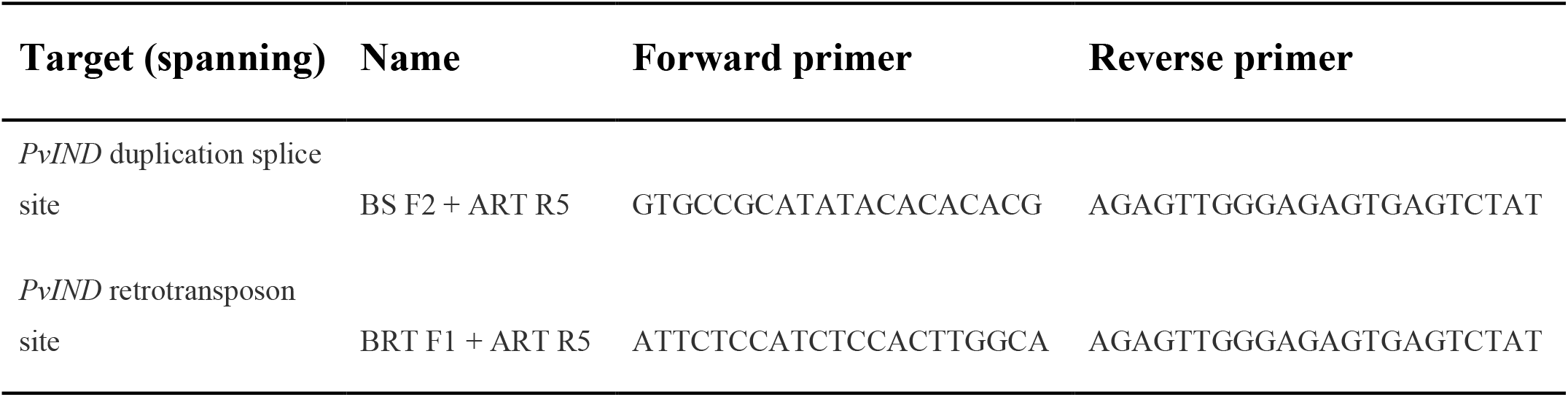
Primers used for characterizing duplication and retrotransposon insertion.

**Supplemental Table S6.**
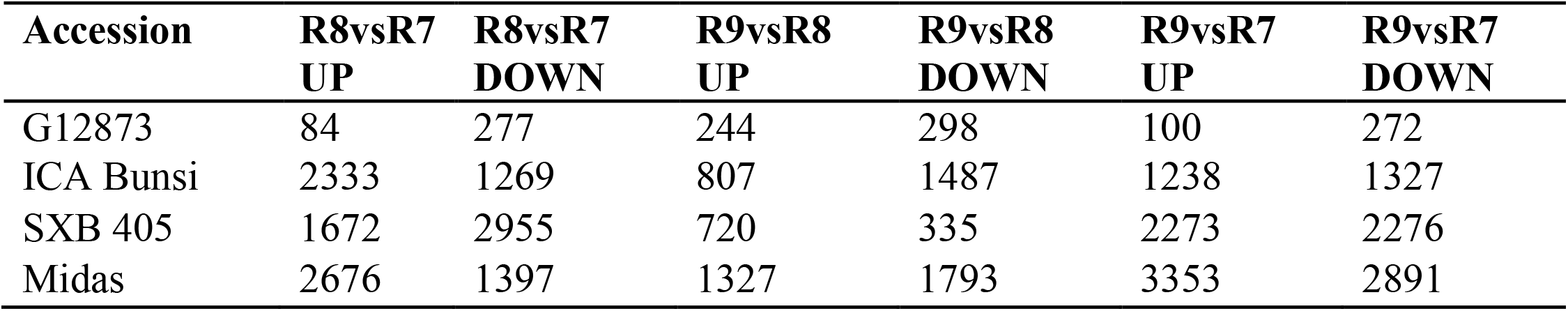
Number of DEGs in the four *P. vulgaris* accessions

**Supplemental Table S7.**
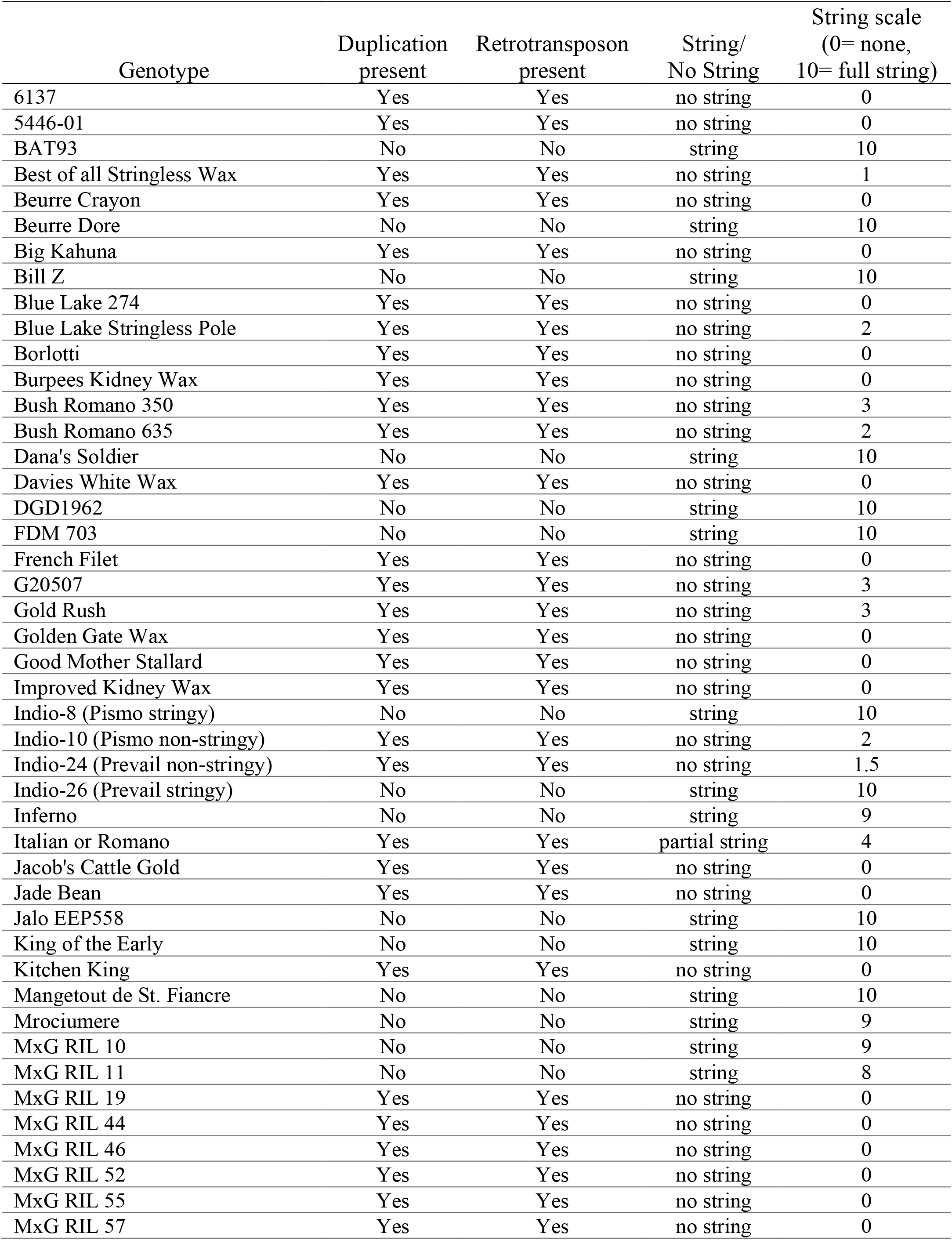
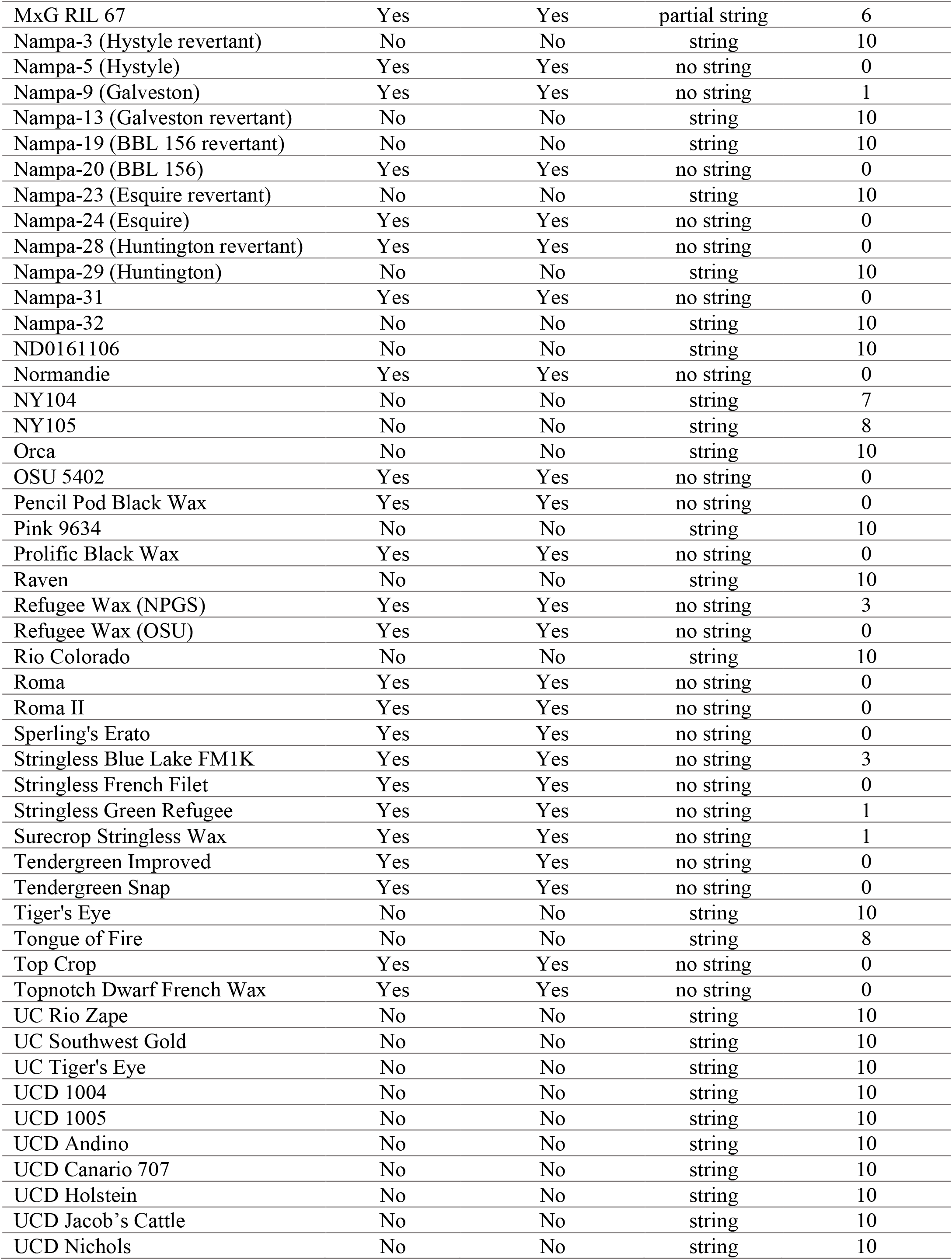
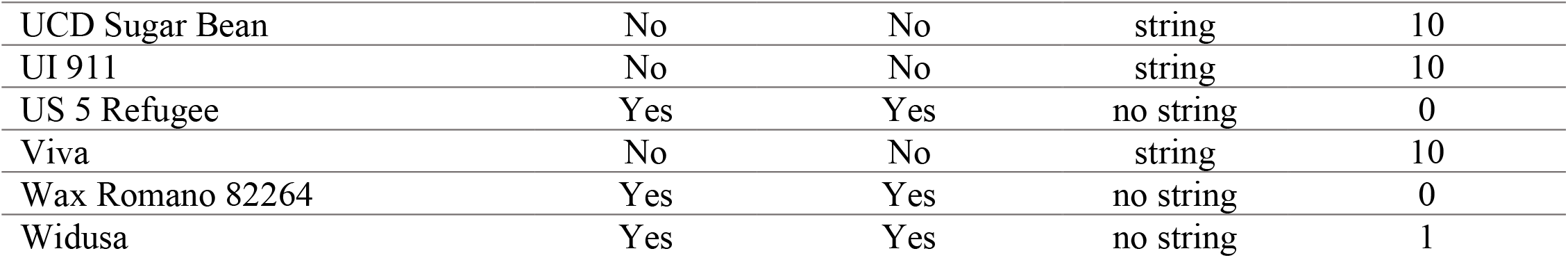
Genotype and phenotype of tested *Phaseolus vulgaris* accessions

